# A pan-cancer atlas of therapeutic T cell targets

**DOI:** 10.1101/2025.01.22.634237

**Authors:** Guangyuan Li, Omar U Guzmán-Bringas, Aman Sharma, Maxence Dellacherie, Palak Sekhri, Rachel Yamin, Dejan Stepec, Maximilien Burq, Ioana Clotea, Ethan Tardio, Aswin Natarajan, Zachary Harpaz, Xinya Liu, David Requena, Darren Taylor, Beatrix M. Ueberheide, Michelle Krogsgaard, C. Russell Y. Cruz, Peter Cimermancic, Mark Yarmarkovich

**Author notes:** Corresponding author: Mark Yarmarkovich, PhD.

## Abstract

T-cell-based immunotherapies have revolutionized cancer treatment, yet only a minority of patients are eligible for these approaches, significantly constrained by the limited knowledge of tumor-specific antigens. Here we present ImmunoVerse, a comprehensive map of T cell targets across 21 cancer types, revealing actionable tumor-specific targets in 89% of tumors analyzed. To define the repertoire of actionable T cell targets, we conducted an exhaustive pan-cancer analysis, integrating data from 7,188 RNA-Seq, 1,771 immunopeptidomes from 512 biological samples and 208 single-cell cancer datasets using novel AI methods, and compared these against 17,384 normal samples covering 51 tissues. Our analysis uncovered 62 viable surface protein targets and 28,446 tumor-specific HLA-presented antigens, deriving from 11 distinct molecular events, across 21 tumor types. Among these, we identified 5,928 previously uncharacterized neoantigens, new tumor self-antigens, peptides derived from tumor-specific cryptic ORFs, tumor-associated microbial targets and a novel splicing-derived PMEL peptide (sPMEL) with enhanced abundance and safety compared to the canonical clinical targets. We successfully expanded sPMEL-specific T cells, validating the therapeutic potential of these targets in functional assays. We highlight 153 promising new tumor targets and experimentally validate 19 targets representing six antigen classes. In addition to being the most comprehensive atlas of targets in scope, ImmunoVerse offers the most extensively annotated resource with key parameters for target selection, providing critical insights for therapeutic prioritization and clinical translation. To catalyze therapeutic development, we released our pan-cancer target atlas through an interactive web portal (https://www.immuno-verse.com) and made the accompanying toolkits available to the scientific community. This work redefines the landscape of therapeutic T cell targets and provides a foundational resource to unlock immunotherapy development across multiple cancers previously considered intractable.

## Introduction

T-cell-based immunotherapies have revolutionized cancer treatment, offering cures to a subset of patients who previously had limited effective therapeutic options^1^. While CAR-T cell therapies have shown remarkable efficacy, particularly in hematologic malignancies such as acute lymphoblastic leukemia and non-Hodgkin lymphoma^2^, their limited success in solid tumors highlights both challenges and therapeutic opportunities. Targeting lineage-restricted proteins shared by malignant and healthy B cells has proven tolerable in blood cancers, as loss of healthy B cells can be managed clinically, but such on-target off-tumor reactivity is unacceptable in solid tumors, where off-tumor reactivity to essential healthy tissues could be catastrophic. This constraint necessitates the identification of CAR targets that offer a substantial therapeutic window. Currently, CAR targeting strategies predominantly focus on membrane proteins, which, while promising, often lack sufficient tumor specificity and have resulted in on-target, off-tumor side effects^3,4^. We and others suggest that the field may be approaching a point of diminishing return in identifying optimal membrane targets for CAR-T therapy^5^, indicating a critical need for innovative targeting strategies that can distinguish between malignant and healthy tissues more effectively.

Human Leukocyte Antigen (HLA) molecules present a snapshot of the cellular proteome on the membrane of all nucleated cells, offering an opportunity to expose a subset of tumor-specific molecular alterations on the cell surface^6^. Targeting peptides present on HLA (pHLA) has achieved curative responses in the clinics through therapies including immune checkpoint inhibitors (ICIs)^7^, adaptive transfer of Tumor Infiltrating Lymphocytes (TILs)^8^, TCR therapies^9^ and recent complete response from neoantigen vaccines^10,11^. Immunotherapies such as ICIs and TILs often rely on high mutational burden, therefore the majority of curative responses to these therapies are applicable only to a limited number of highly mutated tumors. Our group and others have recently demonstrated new approaches, such as peptide-centric (PC)-CARs, that can enable the targeting of any peptide on HLA, including non-immunogenic tumor self peptides, thereby allowing targeting of intracellular antigens in low-to-medium mutational tumors^12^. The prerequisite for developing such immunotherapies is the identification of high-confidence tumor-specific HLA-presented peptides. Tumor-specific targets can be derived from a broad range of genetic aberrations, ideally tumor-dependency genes that underpin the biology of various cancers or other molecular events that promote tumor fitness, yet most of these targets remain uncharacterized. Our group and others have shown targeting through a single tumor-specific pHLA derived from these tumor vulnerabilities that achieve complete anti-tumor responses^13^, underscoring an enormous therapeutic opportunity in extensively characterizing the landscape of targets across tumors.

In the past decade, increasing evidence has revealed actionable HLA-presented epitopes arising from a wide spectrum of tumor-specific genetic aberrations, including canonical protein-coding genes and mutations, alternative splicing^14^, ectopically expressed transposable elements^15^, cryptic ORFs^16^ and post-translational modifications (PTMs)^17^. Many of these findings have been corroborated by immunopeptidomics, the gold standard for empirical characterization of pHLAs by LC-MS/MS proteomics. Standard immunopeptidomic workflows perform spectral matching to a reference database, typically comprised of the canonical proteome, limiting peptide identifications to unaltered self peptides. Our group and others have observed that over 50% of MS/MS spectra remain unannotated when mapped to the database of potential peptides from the canonical proteome – these unannotated spectra represent the “dark matter” of the immunopeptidome^18^. We hypothesize that the molecular instability of cancer results in a multitude of aberrations that can result in potential immunotherapy targets that are not captured using a database of only canonical protein sequences. Hence, uncovering the “dark matter” of the immunopeptidome derived from non-canonical molecular sources will reveal a rich set of additional tumor antigens. While a large number of immunopeptidomic datasets are publicly available, most do not have matched RNA-Seq data, restricting the peptides recovered from these searches to the canonical proteome. Comprehensively identifying all categories of molecular events requires harmonized transcriptomic and immunopeptidomic datasets, multimodal computational workflows, and a well-curated large-scale normal reference to exclude non-tumor-specific molecular events – a challenge that hasn’t been tackled until now. To address these challenges, we developed ImmunoVerse, an exhaustive search engine to comprehensively profile 11 classes of genetic aberrations from RNA-Seq, encompassing all known tumor-specific molecular events. To overcome the limited availability of matched RNA-Seq data, we first generated a histology-matched molecular catalogue of 11 classes of cancer-specific molecular alterations across 21 cancer types utilizing unmatched large datasets from TCGA, TARGET and GTEx. We then utilized this catalogue as a comprehensive search space to interrogate 1,771 immunopeptidomics datasets, revealing a multitude of actionable immunotherapy targets across cancers.

## Result

### ImmunoVerse: A multimodal approach to establish the comprehensive antigen landscape

To comprehensively identify pHLAs derived from diverse molecular events, we developed ImmunoVerse, a multimodal computational pipeline to profile 11 classes of molecular aberrations by integrating tumor transcriptomic and immunopeptidomic data. These 11 molecular sources are known to yield bona fide pHLAs, including canonical protein-coding genes, alternative splicing product, gene fusions, Transposable Element (TE)-chimeric transcripts, self-translating TEs, mutations, intron retentions, tumor-resident microbiome, translatable lncRNAs, pseudogenes and cryptic Open Reading Frames (ORFs). We applied stringent filters for tumor specificity, utilizing up to 17,384 normal tissue samples from Genotype-Tissue Expression (GTEx) to exclude molecular events detected in 51 normal tissues. The retained tumor-specific events were translated into putative HLA peptides, which serves as tumor-specific search spaces to query the histology-matched immunopeptidome datasets. To comprehensively capture all the possible peptide spectrum matches (PSMs), we employed three proteomic search algorithms (MaxQuant^19^, MS2Rescore^20^ and Tesorai Search^21^) to boost the sensitivity. As expected, when comparing to the MaxQuant identifications as baseline, MS2Rescore enhances peptide identification rate by 30% on average, suggesting a benefit to further separating the signal and noise after traditional target-decoy based FDR calculation. Moreover, we applied the newly developed deep learning model Tesorai (trained on over 500 million spectra), which boosted identifications by 60% on average (**Figure S1**). When restricted to high-confidence PSMs (Andromeda score > 70 and Tesorai score > 5), we demonstrated that Tesorai can boost the peptide identification rate by 136% on average (**Figure S2, Table S1**), likely attributed to its incorporation of internal fragment ions in addition to canonical b and y ions, pre-trained scoring function, as well as its single-step FDR estimation. Finally, to minimize false positives, we applied netMHCpan 4.1^22^ to predict HLA binding for all the identified peptides from three different search algorithms^23^ (**Figure 1A,B**). This strategy balanced the improved sensitivity and reduced false discoveries, leading to the identification an abundance of tumor-specific antigens.

**Figure 1.**
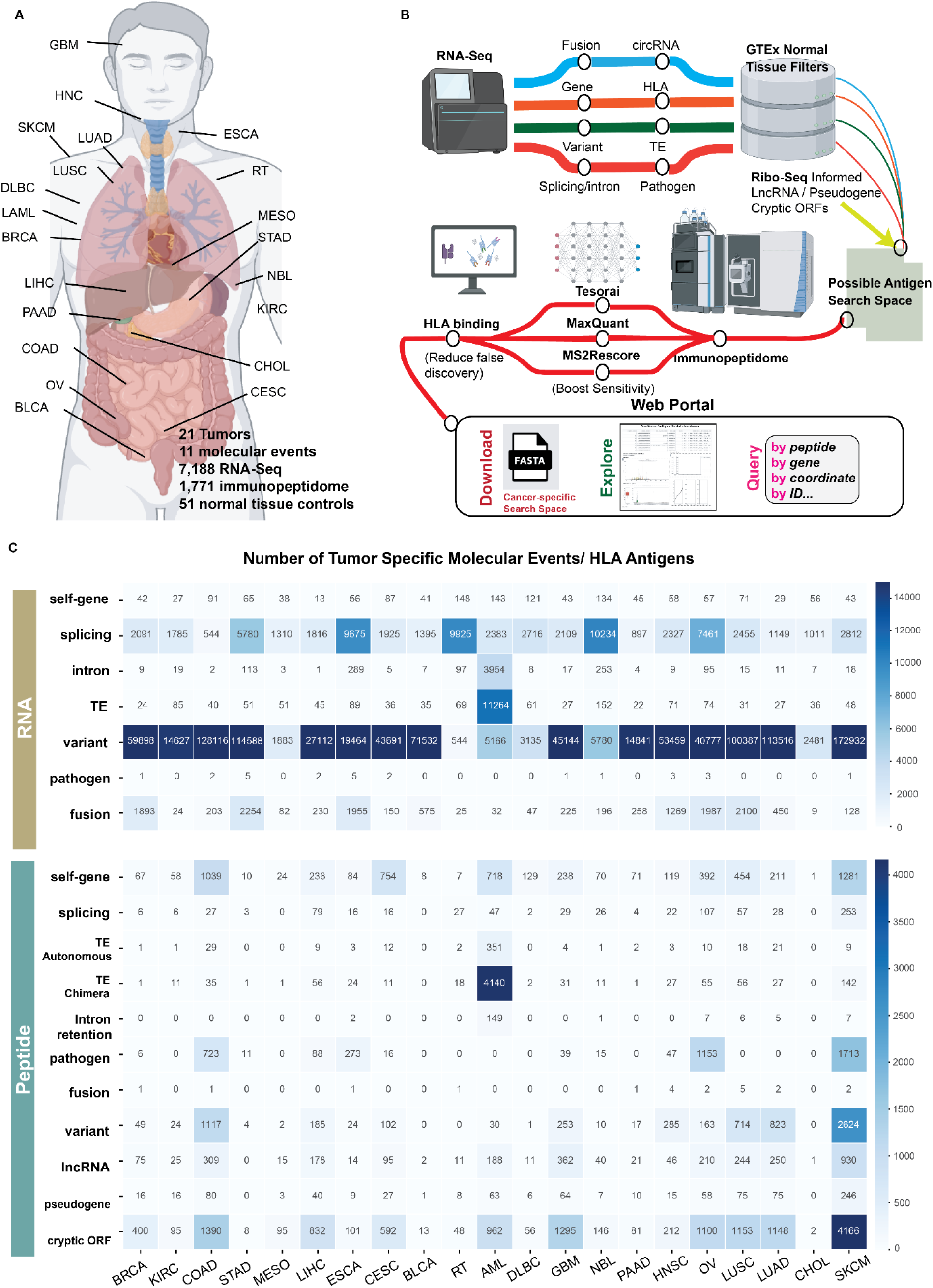
Overview of the analytical pipeline and antigen atlas. (A) Illustration of the cancer types included in this study. Breast Invasive Carcinoma (BRCA), Kidney renal clear cell carcinoma (KIRC), Colon adenocarcinoma (COAD), Stomach adenocarcinoma (STAD), Mesothelioma (MESO), Liver hepatocellular carcinoma (LIHC), Esophageal carcinoma (ESCA), Cervical squamous cell carcinoma and endocervical adenocarcinoma (CESC), Bladder Urothelial Carcinoma (BLCA), Rhabdoid Tumor (RT), Acute Myeloid Leukemia (AML), Diffuse Large B-Cell lymphoma (DLBC), Glioblastoma multiforme (GBM), Neuroblastoma (NBL), Pancreatic adenocarcinoma (PAAD), Head and Neck squamous cell carcinoma (HNSC), Ovarian serous cystadenocarcinoma (OV), Lung squamous cell carcinoma (LUSC), Lung adenocarcinoma (LUAD), Cholangiocarcinoma (CHOL), Skin Cutaneous Melanoma (SKCM). (B) Computational pipeline detailing the steps used to identify, refine, and enhance the detection of tumor-specific antigens, along with a depiction of the functionalities of associated interactive web portals. (C) The distribution of tumor-specific antigens identified across all analyzed cancers, derived from 11 molecular sources, at both the RNA (top) and peptide (bottom) levels.

We sought to apply this pipeline to existing immunopeptidomic datasets, yet found that most available datasets do not have matched RNA-sequencing. To enable broad application of this pipeline to any immunopeptidomic dataset without matched RNA-seq data, we generated a comprehensive catalogue of tumor-specific molecular events to use as a reference database for samples without matched RNA-seq data, enabling us to map the landscape of tumor antigens across histologies. First, we sourced RNA-Seq tumor data from The Cancer Genome Atlas Program (TCGA) and Therapeutically Applicable Research to Generate Effective Treatments (TARGET), and immunopeptidome data from public repositories including Proteomics Identification Database (PRIDE) and Mass Spectrometry Interactive Virtual Environment (MassIVE), along with single cell sequencing data from Cancer Single-Cell Expression map (CancerSCEM) (**Methods, Table S2-3**). Our dataset encompasses 7,188 RNA-Seq, 1,771 immunopeptidome from 512 biological samples and 208 single-cell RNA-Seq datasets across 21 tumor types (**Figure 1A**). Across the 21 tumors analyzed, the number of RNA-Seq samples per cancer ranges from 44 to 1,118, while immunopeptidome samples range from 2 to 352 (**Figure S3A-B**). When incorporating available HLA annotations, AML and SKCM immunopeptidome cohorts exhibit the highest population coverage at 98%. Eleven cancer types achieve over 90% HLA population coverage, while 16 cancers exceed 70% coverage. (**Figure S3C, Table S4**). Although our pipeline is also capable of profiling RNA editing and circular RNAs, detecting circular RNAs typically necessitates specific preparation techniques including depleting ribosomal RNA, enriching for poly(A) negative RNA, or selectively removing linear RNA. Consequently, the poly(A) enriched mRNA protocol predominantly used in TCGA studies leads to underrepresented circular RNA transcripts in our data. RNA editing also has the potential to generate tumor antigens, but it cannot be reliably distinguished from DNA mutations without matched WGS and RNA-Seq data. Therefore, we have not considered these two aberrations in our current pan-cancer analysis.

Applying this strategy, we discovered 28,446 tumor-specific pHLAs across 21 tumors deriving from all 11 classes of aberrations. Amongst the identified somatic mutations, high mutation burden tumors such as melanoma and lung squamous cell carcinoma exhibited the highest counts, followed by colon and stomach cancers, which are often characterized by microsatellite instability. Conversely, immunologically cold tumors such as mesothelioma and neuroblastoma show the lowest number of mutations. Strikingly, Acute Myeloid Leukemia (AML) shows the highest prevalence of tumor-specific transposable elements (n=11,264) and intron retention events (n=3,954). It has been reported that AML are characterized by widespread intron retention events^24^ and endogenous retroviruses could serve as genetic enhancers in AML that drive tumorigenesis^25^, suggesting a potential interplay between epigenetic changes, endogenous retroviruses, and alterations in the splicing landscape (**Figure 1C**). When calculating the number of tumor-specific pHLAs across cancers, melanoma and AML have the highest mutation-derived and TE-derived antigens, respectively, as expected based on their high burden of underlying genetic aberrations. However, discrepancies are observed in other cancers like neuroblastoma, which, despite possessing a high number of tumor-specific alternative splicing events detected from RNA-Seq, yields fewer pHLAs derived from alternative splicing (**Figure 1C**), demonstrating a discordance between transcripts and their representation on HLA. Notably, the low level of antigen identification in stomach cancer and cholangiocarcinoma can be attributed to limited sample availability in the immunopeptidome data (STAD: 2, CHOL: 5) and suboptimal HLA population coverage (STAD: 58%, CHOL: 26%), underscoring the need for additional immunopeptidomic studies for such cancer types. Taken together, our work provides a comprehensive and actionable landscape for exploring tumor antigens, spanning a variety of molecular events across tumors.

The identified tumor antigens are accessible through our web portal (https://www.immuno-verse.com/). Moreover, we have released our pipeline as a code-free cloud-based pipeline through Cancer Genomics Cloud (CGC), allowing users to process their own immunopeptidomic and/or RNA-seq data through ImmunoVerse. We have also made the putative antigen search space available for each tumor type, allowing users to conduct comprehensive immunopeptidomic searches with either their RNA-Seq data and/or immunopeptidomic data. Taken together, the availability of these resources to the research community substantially expands accessibility.

### Membrane proteins provide limited new therapeutic opportunities

We first evaluated the landscape of surface proteins suitable for conventional CAR-T therapy, applying the identical filtering stringency used for pHLA targets to all annotated human cell membrane proteins with exposed extracellular domains. This unbiased analysis identified 89 membrane protein candidates across tumors with clear therapeutic windows by comparing their gene expression in tumors to all normal human tissues (**Methods**). As expected, this list includes TNFRSF17/BCMA and CD19, the only two FDA-approved CAR-T targets to date, along with fourteen additional targets currently being investigated in clinical trials (CLEC12A, DLL3, ULBP2, MUC16, FLT3, MSLN, CD70, CDH17, FOLR1, IL1RAP, GPC2, CLDN6, CEACAM5, TNFRSF13B). Further literature review shows 11 additional targets that have established CARs and are actively being tested in different preclinical models. Extending from previously reported targets, our analysis also highlighted 62 previously unrecognized antigens with favorable therapeutic windows (**Figure 2, Table S5**), suggesting their suitability as novel candidates for conventional CAR-T therapeutic development and warranting further investigation. Notably, several clinical and preclinical CAR targets, including ERBB2/HER2, EPCAM and MUC1, did not pass our filtering, consistent with reported preclinical and clinical toxicities or a lack of a therapeutic window for effective targeting^26–28^. CAR targets such as GD2, a glycosphingolipid, are not identifiable through our RNA-based analysis. While previous CAR-T target discovery efforts have only focused on a limited subset of genes supported by protein staining data and often considered the tumor specificity of currently approved CAR-T targets as an acceptable benchmark^4^, our analysis expands the search to all membrane proteins and applies more stringent specificity criteria. This analysis both reveals new membrane proteins that can serve as potential conventional CAR targets and underscores the need for the identification of additional tumor antigens beyond canonical membrane proteins.

**Figure 2.**
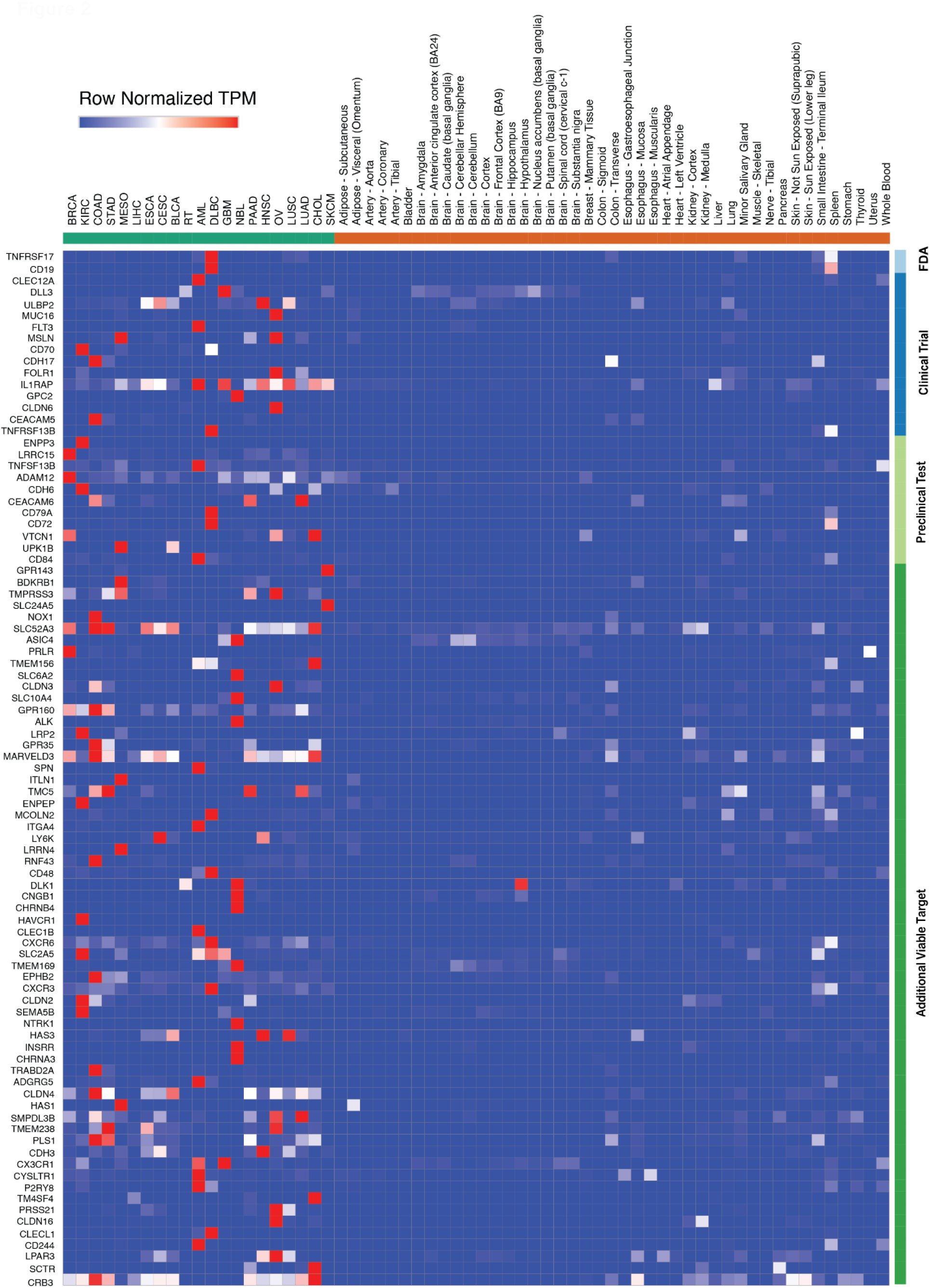
Suitable tumor-specific surface proteins for conventional CAR-T therapy. Surface targets (n=89) are categorized and color-coded based on their development status including FDA-approved, in clinical trials, in preclinical testing, or classified as additional viable candidates. The heatmap displays row-normalized Transcript Per Million (TPM) values for each gene. Selection criteria (**Methods**) requires genes to be highly expressed in tumors, exhibit an appreciable therapeutic window relative to normal tissues, show low expression in essential normal tissues, and, where applicable, be significantly overexpressed compared to matched normal controls. To ensure specificity, we manually inspected all retained targets and excluded those with appreciable expression in essential normal tissues (blacklist genes; **Table S5**). Only targets that passed manual review are highlighted.

### Self-antigens represent a major source of PC-CAR targets

In our recent work, we established the use of PC-CARs to target low-to-medium mutational tumors, enabling potent and specific targeting of oncogene-derived peptides abundantly presented on HLA molecules^12^. Our initial efforts focused on neuroblastoma, where we targeted the developmental transcription factor and master regulator in neuroblastoma tumor maintenance PHOX2B. This factor is silenced in healthy differentiated tissues but is co-opted by tumor cells to drive tumorigenicity. PC-CARs engineered against PHOX2B demonstrate complete elimination of tumors in preclinical models^29^ and are now in a Phase I clinical trial (NCT07007117). These results underscore the therapeutic potential of uncovering a new tumor-specific target, we therefore sought to expand this therapeutic strategy to a broader range of cancer patients.

To map the landscape of actionable epitopes for T-cell-based therapies, we scored canonical proteins in our immunopeptidomic datasets by (1) abundance, (2) recurrence, (3) absence in normal tissue RNA and immunopeptidomes, (4) HLA binding affinity, (5) essentiality in maintaining tumor progression, (6) homogeneity as determined by single cell coverage, and (7) confidence level of mass spectrometry-based identification. We identified 4,522 tumor-specific peptides derived from 292 genes that are highly-enriched in tumors without appreciable level of expression in essential normal tissues, drastically expanding the current target repertoire (**Table S5**). Expression profiling revealed distinct clusters of genes exclusive to specific cancers (**Figure 3A**). Notably, cancers such as melanoma, AML, and ovarian cancer harbor uniquely expressed genes, including well-characterized immunotherapy targets like PRAME, MPO and MUC16. Consistent with this, clustering of 1,771 immunopeptidome datasets based on peptide abundance reveals that these cancer types are more readily separable than others, owing to their distinct tumor-specific antigens (**Figure 3B**). Surprisingly, we showed the immunopeptidome landscape, while less pronounced than transcriptome, can still enable discrimination across major cancer types and suggest its potential utility for diagnostic applications using serum immunopeptidomics.

**Figure 3.**
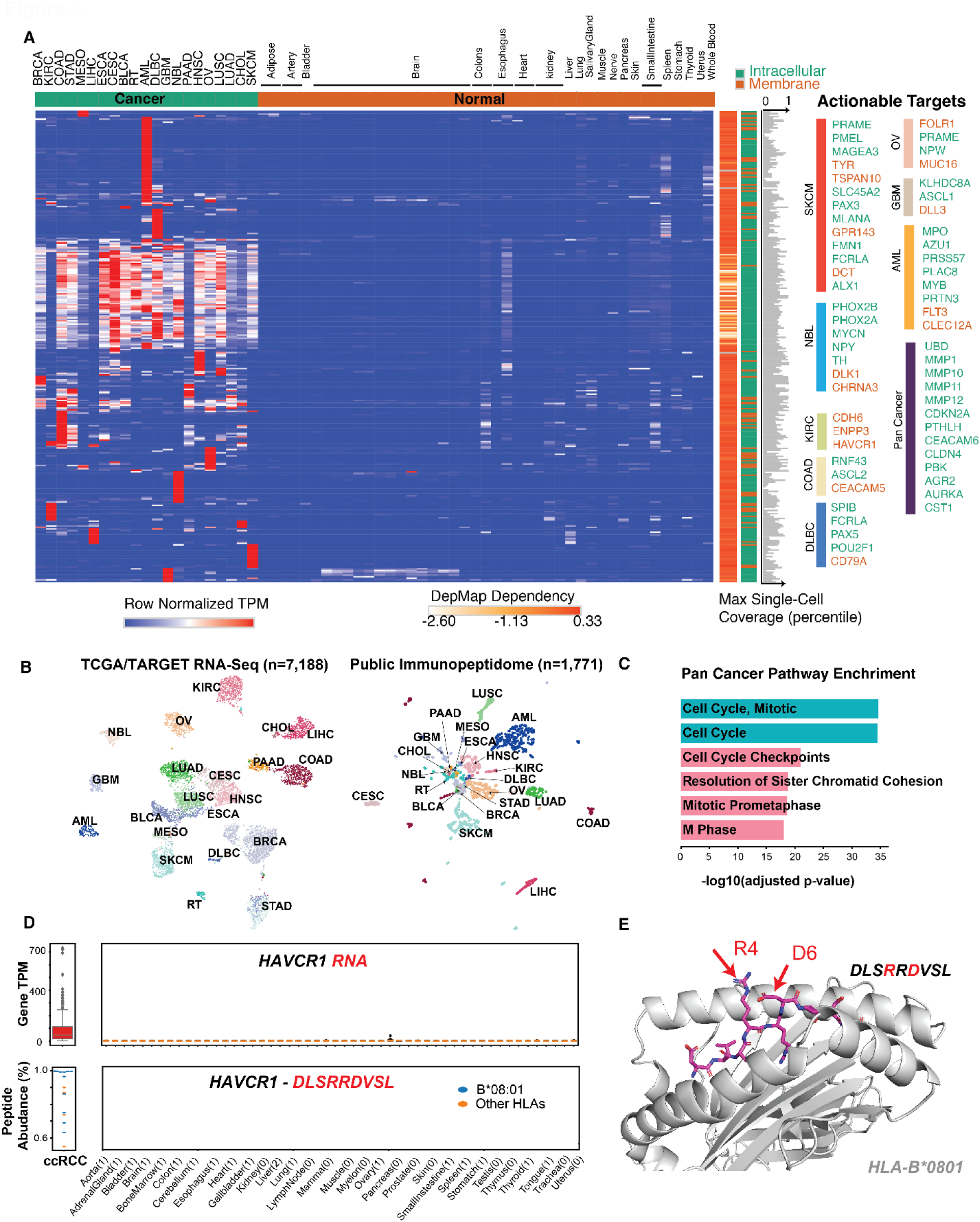
Landscape of pHLA from tumor specific self-antigen. (A) Gene expression of 292 tumor specific genes that can give rise to tumor-specific pHLA antigens, across 21 tumors and normal tissues. Genes are annotated by whether is membrane or intracellular proteins, along with the DepMap reported dependency score (median across all evaluated cell lines) and homogeneity represented as max single cell RNA-Seq coverage (shown in percentile based on ranking all detected genes). Gene expression is depicted by median transcripts per kilobase million (TPM) in each cancer or tissue. (B) Uniform Manifold Approximation and Projection (UMAP) of 7,188 RNA-Seq samples based on the transcript-per-million (TPM) values of 13,033 genes known to give rise to peptides detected in the immunopeptidome (left), along with UMAP of 1,771 public immunopeptidome datasets based on the median peptide abundance attributed to each parental gene (right). (C) Enriched gene ontology and pathway associated with 67 clustered pan-cancer targets. (D) HAVCR1 shows high differential expression by RNA (top) and peptide abundance (bottom) when comparing between clear cell Renal Cell Carcinoma (ccRCC) and normal tissues. The number of normal immunopeptidome samples with HLA-B*08:01 is indicated in parentheses for each tissue type. (E) Alphafold2 modeling of HAVCR1 peptide and cognate HLA-B*0801 complex, with exposed two charged residues (R4, D6) highlighted.

We identified a cluster of peptides derived from genes involved in cell cycle (CDC6 and CDC45), invasion and extracellular matrix remodeling (PTHLH, MMP1, MMP10), which are broadly expressed across multiple tumors (**Figure 3A,C**), highlighting shared pathways of tumor progression that are exposed on HLA across tumors. Although cell therapies targeting these common vulnerabilities alone may affect normal proliferative cells, given the role of these genes and broad upregulation in cancers, incorporating them into AND-gated strategies^30,31^ with other antigens could represent a viable therapeutic strategy.

To determine how many cancer patients could potentially benefit from pHLA-centric therapies, including PC-CARs and engineered TCRs, we analyzed gene expression profiles across all cancer patients. Our findings reveal that all of the cancers in our study harbor at least one tumor-specific gene capable of generating tumor self-antigens. Even when taking into account empirically-evidenced peptides and HLA restrictions, more than 99% patients can potentially benefit from PC-CARs for 14 out of 21 cancer types, demonstrating the broad potential of these therapies. However, some cancers, like mesothelioma and rhabdoid tumors, showed lower patient coverage, underscoring the need for expanded immunopeptidomic profiling (**Figure S3D**). Highlighting an example, we identified a self-antigen DLSRRDVSL originating from tumor-specific gene HAVCR1 and present by a common allele HLA-B*08:01, covering over 15% of the U.S. population. While HAVCR1 is a potential surface protein target, it is reported to shed its ectodomain from the cell surface, providing a potential evasion mechanism to conventional CARs^32^. Thus, we suggest that targeting HAVCR1 presented on HLA using PC-CARs provides an alternative therapeutic strategy. The HAVCR1 peptide is absent in normal immunopeptidomic datasets and exhibits a favorable therapeutic window at both RNA and peptide levels (**Figure 3D**). Furthermore, AlphaFold2 modeling predicts the binding to the cognate HLA and several key biophysical features that offer a favorable surface for targeting with two exposed charged residues accessible for salt bridge formation (**Figure 3E**). We validated this antigen using synthetic peptide and compared the resulting LC-MS/MS spectra, demonstrating complete concordance across all b and y ions (**Figure S4A,B**). Additional examples of tumor self-antigens include lineage-restricted markers such as POU2AF1 in diffuse large B-cell lymphoma, oncoproteins from the CEACAM family in colon cancer, and known differentiation antigens like MLANA/MART1 in melanoma (**Figure 3A, Table S5**). The full catalogue of tumor-specific self-antigens is available through our web portal, which we encourage investigators to explore. Taken together, our comprehensive pan-cancer analysis underscores the substantial potential of targeting HLA-presented self-antigens across a wide range of cancers using PC-CARs and other immunotherapies.

### ImmunoVerse reveals numerous empirically-evidenced mutation-derived neoantigens

While neoantigens arising from somatic mutations and gene fusions offer inherent tumor specificity that has been successfully leveraged in clinical trials, these peptides are typically selected based on computational predictions that incorporate gene expression and HLA binding affinity. Indeed, it has been reported that only 2.4% of predicted strong pHLA binders are detected within the immunopeptidome^33^ and only 0.5-2.3% of predicted antigens demonstrate immunogenicity across multiple types of cancers^34^. Therefore, peptide vaccine cocktails based on computational predictions contain a high proportion of peptides that may not be presented by tumors.

We reasoned that an optimal neoantigen target for cancer vaccine and TILs expansion should be supported by empirical immunopeptidome evidence, confirming both its existence and abundance in tumors. To quantify the prevalence of such optimal neoantigens and prioritize them, we systematically searched all SNV- and INDEL-derived mutation peptides across our immunopeptidomic cohorts, encompassing both public and private antigens. This analysis revealed 6,403 mutation-derived neoantigens with direct immunopeptidome evidence. The contribution of mutation-derived neoantigens toward all non-canonical tumor-specific pHLAs varied across cancer types, with head and neck cancer exhibiting the highest proportion (46.4%), and an average contribution of 14.6% amongst all non-canonical pHLAs (**Figure 4A**). Specifically, skin cutaneous melanoma (SKCM), colon cancer and lung adenocarcinoma exhibited the highest numbers of mutation-derived neoantigens, with counts of 2,624, 1,117, and 823, respectively, consistent with reported mutational frequencies at genetic level^35^ (**Figure 4B**). Missense mutations account for 87% of all neoantigens, followed by frameshift mutations at 11.7%. Neoantigens derived from in-frame insertions and deletions are rarely detected, contributing only 0.005%. (**Figure 4C**). Comparing our findings to the prediction-based pan-cancer neoantigen database TSNAdb^36^, we find that only 475 of their reported neoantigens had immunopeptidome evidence within our cohort (**Figure 4D**). These accounted for only 0.001% of the previous total prediction neoantigen list, underscoring the importance of empirical characterization in therapeutic target discovery and selection.

**Figure 4.**
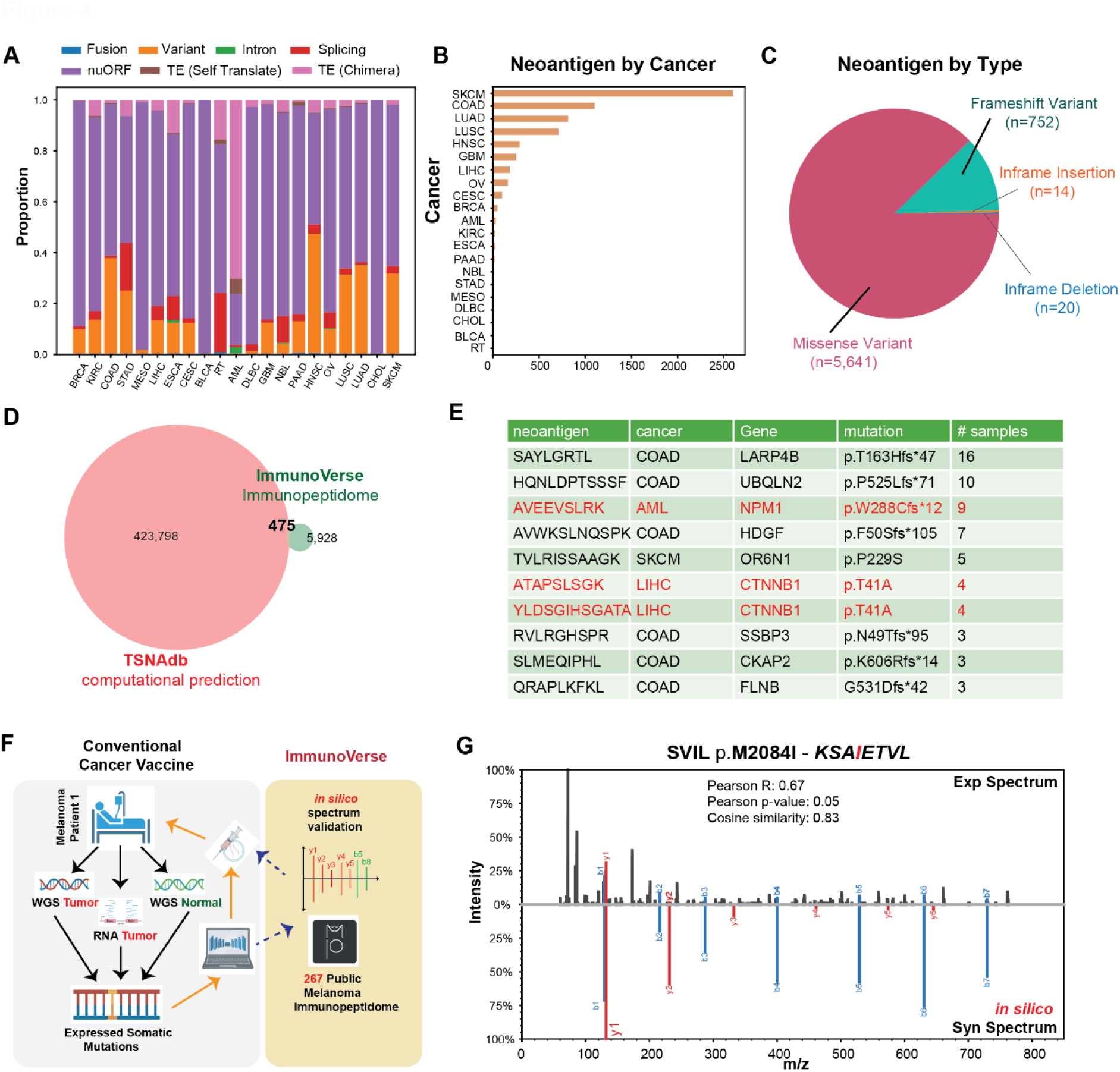
Landscape of mutation-derived neoantigens across cancers. (A) Contribution of diverse non-canonical pHLAs identified from immunopeptidome data across different cancer types. (B) Number of experimentally evidenced neoantigens detected per cancer type. (C) Distribution of evidenced neoantigens categorized by mutation type. (D) Comparison between computationally predicted neoantigens collected through TSNAdb database and those confirmed by immunopeptidomics through the study. (E) Representative list of public neoantigens supported by immunopeptidome evidence, neoantigens derived from tumor driver genes are highlighted in red. (F) Illustration of integrating ImmunoVerse pipeline and curated immunopeptidome datasets to prioritize neoantigens in clinical setting. Specifically, conventional pipeline (grey background) processes Whole Genome Sequencing (WGS) data between tumor lesions and matched normal tissue, along with tumor RNA-Seq to identify expressed neoantigens. Subsequently, expressed neoantigens are only subject to computational predictions. The added module (yellow background) proposes a direct search of expressed neoantigens against public immunopeptidome data followed by *in silico* prediction of the neoantigen spectra to evaluate the confidence of identifications before resuming the patient-specific binding and immunogenicity evaluation. (G) One representative neoantigen (KSAIETVL) evidenced through public melanoma immunopeptidome and is evaluated on the fly with high confidence scores.

Among the newly identified immunopeptidome-evidenced neoantigens, we highlight shared neoantigens in colon cancer derived from frameshift mutations in LARP4B, as well as a missense mutation, CTNNB1 T41A, a driver gene in liver cancer. We further confirmed their validity by comparing the MS/MS spectra to synthetic peptides (**Figure 4E**, **Figure S4C,D**). Importantly, our analysis also confirmed several well-characterized shared neoantigens, including AVEEVSLRK derived from an NPM1 frameshift mutation in AML, SLMEQIPHL from a CKAP2 frameshift mutation in colon cancer, and 37 private neoantigens now deposited in the IEDB database. Furthermore, we identified novel neoantigens derived from gene fusions, which, although typically patient-specific, may represent promising immunotherapeutic targets. These include NCOA7-TPD52L1 in breast cancer, GNG5-NMI in colon cancer and FRAS-MRPL1 in lung squamous cell carcinoma (**Table S6**). Taken together, our pan-cancer molecular catalogue provides a valuable resource for identifying both previously reported and novel neoantigens. The direct detection of these antigens in the immunopeptidome strengthens the case for prioritizing them in the development of cancer vaccines or adoptive TIL therapies.

Given the ability to detect both shared and private neoantigens using histology-matched public immunopeptidome data, we sought to directly integrate our tool into the selection of peptides for cancer vaccines. Specifically, we hypothesized that somatic mutations identified through whole-genome and RNA sequencing could be directly searched against our curated immunopeptidome database, allowing high-confidence matches to be prioritized for vaccine development based on their likelihood of natural presentation in tumors (**Figure 4F**). As a proof of concept, we analyzed RNA-Seq and paired whole-genome sequencing data from one melanoma patient. Amongst the identified 537 expressed somatic mutations from WGS and RNA-Seq, we subsequently searched these against 267 public melanoma immunopeptidomic datasets, identifying eight neoantigens with supporting evidence. While orthogonal validation using synthetic peptides is possible, to fit the constrained time-scales of vaccine manufacturing, we incorporated a feature into ImmunoVerse that can computationally predict the synthetic spectrum and evaluate the confidence of each identification (**Figure 4G**). Further prediction of HLA binding affinity and immunogenicity revealed that one of the neoantigens, FLRTFLGGRF, derived from the FLYWCH1 S604F mutation, is predicted to bind the patient’s HLA-B*27:02 allele and demonstrates high predicted immunogenicity (DeepImmuno^37^ score = 0.99), suggesting that this peptide would be more likely to be presented than peptides selected solely based on computational predictions and induce a T cell response. Moreover, the entire workflow takes less than four hours to process, allowing for seamless integration into current cancer vaccine workflows while meeting rapid fast turnaround time. We anticipate that incorporating empirically supported neoantigens into vaccine and TIL therapy designs will enhance their precision and clinical efficacy

### Cryptic ORFs and alternative splicing product yield actionable targets

Recent studies have indicated that cryptic open reading frames (ORFs), defined as previously unannotated ORFs originating from untranslated regions or overlapping with canonical ORFs, can be dependencies for tumor growth^38^ and generate actionable targets across various cancers, potentially yielding targets that are more widely shared across patients^39–41^. In line with the previous report, our analysis revealed dominant cryptic-ORF-derived antigens contributions, accounting for approximately 72.8% of the non-canonical repertoire on average (**Figure 4A**). Particularly, the cryptic ORFs give rise to a rich source of tumor-specific antigens that are often recurrent across cancers (**Figure 5A, Table S7**). To validate the translational activity of these cryptic ORFs, we utilized ribosomal profiling (Ribo-Seq) on eight matched neuroblastoma patient-derived xenograft (PDX) models (**Figure S5A**). This analysis confirms the presence of ribosomes on 37% of these ORF-derived pHLAs, indicating that they are actively translated and are able to generate bona fide peptides on HLA (**Figure S5B, S6**). One notable example is the peptide STIPVLSGY from the downstream untranslated region (UTR) of the TMEM203 gene, which was confirmed by both LC-MS/MS of the synthetic peptide and refolding experiments (**Figure S5C-E**). We also identified this tumor-specific antigen in cervical cancer, lung cancer, glioblastoma and melanoma, underscoring its potential as a pan-cancer target.

**Figure 5.**
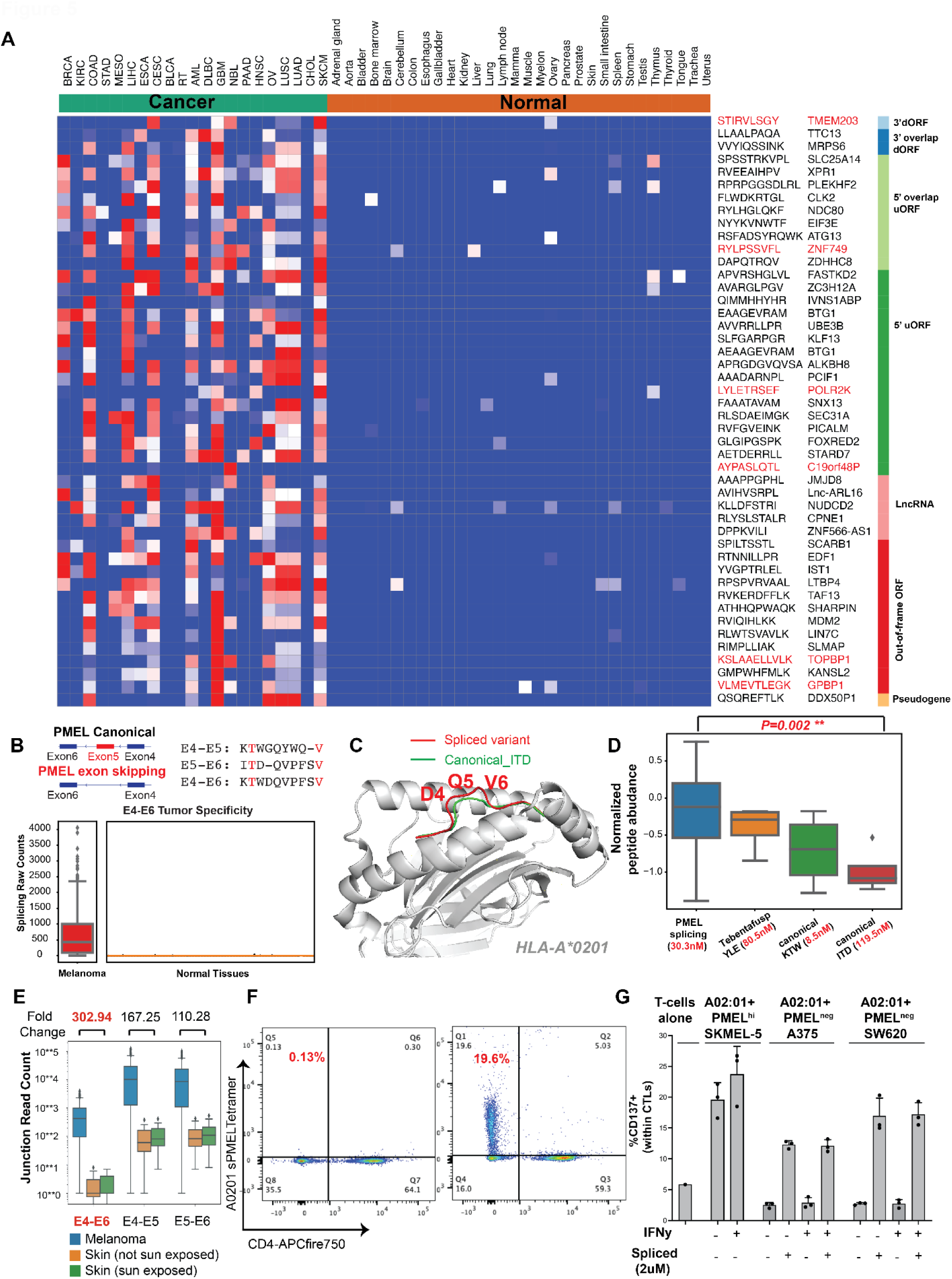
Cryptic ORF and alternative splicing contribute to antigen repertoire. (A) Row normalized peptide abundance between tumor and normal immunopeptidome of representative cryptic ORF derived antigens selected by their recurrence, categories and Ribo-Seq support, annotated by the parental gene and categories. Antigens supported by Ribo-Seq in neuroblastoma is highlighted in red (B) exon skipping of exon 5 of PMEL results in a splice variant compared to two canonical antigens (top) and the supporting RNA-Seq reads of this exon skipping in melanoma versus normal tissues (bottom). (C) Alphafold2 structural modeling of the PMEL splice variant backbone with its cognate HLA-A*02:01 allele, compared to the canonical antigen ITDQVPFSV with distinct TCR-recognizable residues are highlighted. (D) Normalized peptide abundance of the PMEL spliced variant, tebentafusp antigen and two canonical counterparts, the predicted binding affinity of A0201 were annotated for each peptide. (E) Therapeutic windows on RNA junction level between spliced and canonical junctions in cutaneous melanoma. The fold change is derived between melanoma and sun-exposed skin (F) Representative flow plots showing HLA-A*02:01 pMHC tetramer binding to primary T-cells expanded against spliced PMEL peptide KTWDQVPFSV (gated on CD3+ T-cells). (G) Quantification of CD137 expression shows spliced PMEL-dependent activation in expanded T-cells.

In examining splicing-derived antigens, we identified a rich source of tumor-specific splicing junctions that were distinct from those in normal tissues and clustered into cancer-specific events (**Figure S6D, S7A, Table S7**). One particularly notable finding was the recurrent detection of a splicing variant from PMEL exon 5 skipping, which results in a distinct antigen that retains both HLA-A2 anchor sites shared by two other known PMEL epitopes derived from the canonical protein (**Figure 5B**). To confirm the validity of our identification, we compared its MS/MS spectrum to a synthetic peptide and performed a refolding assay to verify its binding to HLA-A*02:01. (**Figure S7B,C**). We propose that this novel epitope can be uniquely recognized by TCRs as compared to the canonical PEML epitopes, highlighting a distinct geometry using structural modeling and showing that two TCR clones against canonical PMEL (Exon5-Exon6) do not cross-react against the non-canonical peptide (**Figure 5C, Figure S7D**). Despite lower transcript abundance, this splicing variant is more abundantly presented on HLA compared to its canonical counterparts KTWGQYWQV, ITDQVPFSV and YLEPGPVTA, a FDA-approved ImmTac target in uveal melanoma, possibly due to enhanced degradation of short-lived unstable splicing products (**Figure 5D, Figure S7E**). Similarly, the spliced antigen is more broadly presented and detectable in melanoma patients compared to other three antigens (**Table S7**). Vitiligo is a known side effect of current melanoma immunotherapy, likely due to the presence of cancer specific targets in normal melanocytes^42^. In our safety assessment of the spliced junction, we observed a superior therapeutic window in both cutaneous and uveal melanoma with negligible expression of the spliced junction in normal skin (**Figure 5E, Figure S8A**). Notably, the spliced antigen exhibits an elevated therapeutic window in cutaneous melanoma compared to uveal melanoma (**Figure S8B**).

To test the capacity to expand T cells against spliced PMEL (sPMEL) antigen and their ability to recognize melanoma through the splicing junction, we expanded HLA-matched T cells from normal donors and co-cultured them with monocyte-derived dendritic cells, resulting in 52.9% of antigen-specific cytotoxic T-cells following 3 rounds of expansion (**Figure 5F, Figure S8C,D**). Co-culturing expanded T cells with HLA-A*02:01 but PMEL-negative lines resulted in activation only upon the addition of sPMEL peptides, demonstrating the ability of expanded cells to activate in an antigen-specific manner. To test for the ability of expanded T cells to recognize tumor cells through endogenously presented sPMEL, we co-cultured expanded T cells with HLA-A*02:01, PMEL-positive SKMEL5 cells, demonstrating potent T cell activation (**Figure 5G, Figure S8E**). Given its likely superior therapeutic properties compared to the canonical PMEL antigen currently targeted in the clinic in conjunction with the feasibility of isolating sPMEL-specific TCR clones, our findings support further exploration of sPMEL-based vaccines, TCRs, T cell engagers, and PC-CARs.

Additionally, as one subset of aberrant splicing variant, we identified pHLAs derived from retained ATP10A intron 8, SUSD1 intron 19 and KLHL5 intron 11, all specific to AML (**Table S7**). The high prevalence of intron retention-derived antigens in AML aligns with previous reports of abundant intron retention aberrations in AML, which have been correlated with mutations in epigenetic modifiers such as DNMT3A^24^ and splicing factors such as SRSF2^43^. Collectively, our findings reveal that tumor-specific antigens derived from alternative splicing events and cryptic open reading frames contribute additional promising targets across various cancers, with significant potential in expanding immunotherapies.

### Tumor resident pathogens provide an unexpectedly rich set of potential therapeutic targets

Viral infections including Hepatitis B Virus (HBV) and Human Papillomavirus (HPV) are well-documented oncogenic viruses directly linked to liver and cervical cancer tumorigenesis, respectively. In HBV and HPV, the integration of viral DNA into the host genome and the direct encoding of transforming proteins have been shown to disrupt cellular regulatory networks, inducing oncogenesis through the activation of oncogenes and inhibition of tumor suppressor genes^44^. Prophylactic vaccinations against these viruses have proven effective in reducing cancer incidence, suggesting that the immune system is able to clear infected precancerous cells through viral antigens^45,46^. We hypothesize that tumor-resident viral pathogens not only contribute to oncogenesis but may also alter the immunopeptidomic landscape by presenting novel antigens, which could reveal new therapeutic targets. To confirm our ability to identify pathogen-derived peptides in tumors, our initial studies focused on the immunopeptidome of liver and cervical cancers known to harbor HBV and HPV infections (**Figure 6A**). Notably, we identified several known viral peptides also residing in primary tumor tissues, such as FLLTRILTI from the HBV large surface antigen, and YMLDLQPET from the HPV protein E7. A comparative analysis with the Immune Epitope Database (IEDB) revealed that 40% of the HBV peptides detected in primary liver tumors were previously documented as immunogenic, highlighting their potential utility as therapeutic vaccine targets (**Table S8**). Building on these observations, we further explored other common oncogenic viruses that are linked to cancers. While cytomegalovirus (CMV) infection has been implicated in multiple cancer, and its DNA and protein have been detected in cancers such as neuroblastoma, colon cancer and glioblastoma, it is currently unclear whether CMV infection can generate additional pHLAs suitable for therapeutic development. Strikingly, our pan-cancer analysis identified 187 CMV-derived antigens across six different tumor types (**Table S8**). Cross-referencing the CMV peptides identified in primary tumors with the IEDB database revealed five peptides that have been previously documented in the setting of viral infection. In particular, we identified the CMV-derived peptide LLDGVTVSL from the essential UL49 protein, a key regulator of CMV gene transcription^47^, which was recurrently detected in seven cancers (ovarian cancer, breast cancer, colon cancer, liver cancer, neuroblastoma, glioblastoma, cervical cancer) and presented by the common HLA-A*02:01 allele (**Figure 6B**). To the best of our knowledge this is the first time these CMV antigens have been reported in tumors, offering a potential new therapeutic avenue using various immunotherapeutic modalities.

**Figure 6.**
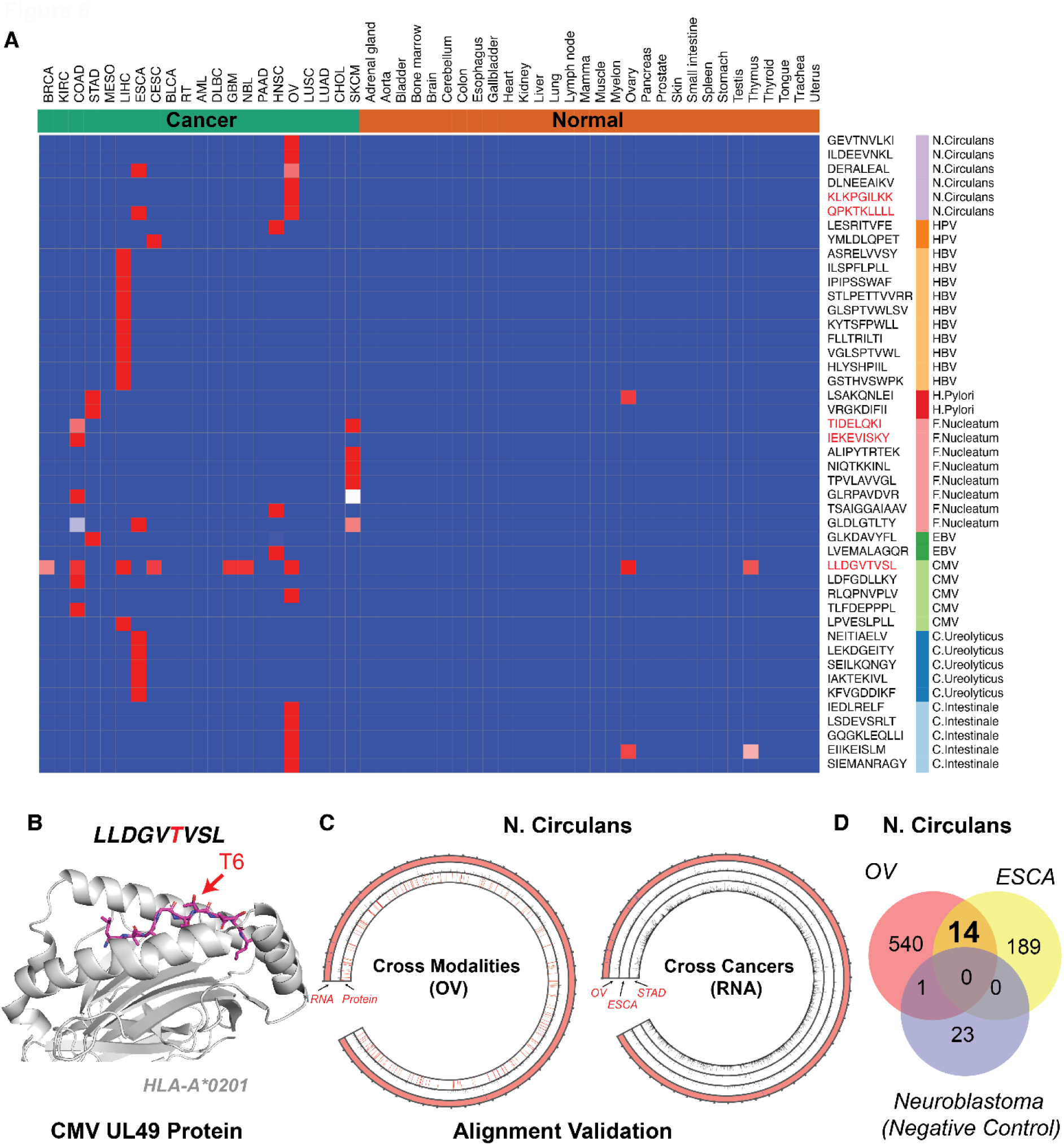
Tumor-specific microbial antigens detected across cancer types. (A) Tumor-specific antigens derived from diverse tumor-resident microbes, selected by immunopeptidome evidence and identification confidence. (B) AlphaFold2 structural modeling of the CMV-derived peptide LLDGVTVSL bound to HLA-A*02:01, identified across multiple cancer types. (C) Circos plot illustrating ovarian tumor RNA-Seq reads and translated protein fragments unambiguously mapped to the *N. circulans* reference genome and proteome (left), along with RNA-Seq reads uniquely mapped to the *N. circulans* genome across ovarian, esophageal, and stomach cancers (right). Blue ticks indicate annotated 16S rRNA regions. (D) Venn diagram showing the overlap of *N. circulans*-derived peptides detected in ovarian cancer, esophageal cancer, and neuroblastoma (used as a negative control).

In addition to oncogenic viruses, we sought to investigate the presence of additional microbes, including bacteria and fungi, that could result in novel antigens. Although it is possible to predict the presence of pathogens from bulk transcriptome data, the existence of a tumor-specific microbiome remains contentious due to the complexity of distinguishing genuine microbial sequences from host-derived or environmental contaminants^48^. This ambiguity is exemplified by the fact that many unmapped reads can be reclassified using the latest reference human genome assembly or may simply be artifacts of contamination, leading to potential false discoveries^49^. To address these discrepancies, we established a stringent framework to re-classify unmapped reads into various species including fungi and bacteria along with human reference, assigning a read to a specific species only if 90% of its kmers (short, fixed-length nucleotide sequences), could be unambiguously mapped to that species. Additionally, we excluded species detected in normal tissue to ensure tumor specificity and corroborated our findings with previously published studies^50^.

Our analysis successfully identified several recurrent strains at the RNA level that also generate peptides in the primary tumor immunopeptidomes, including *H. pylori*, a pathogen associated with gastric cancer tumorigenesis, and *F. nucleatum*, a well-described pathogen implicated in colon cancer and melanoma^51,52^. In addition to previously reported *F. nucleatum*-derived peptides in colon cancer and melanoma, we also identify their presence in head and neck as well as esophageal cancers, consistent with prior reports of *F. nucleatum* colonization in these tumor types^53^ (**Figure 6A**). Aside from these previously described strains, we detected numerous novel strains including *N. circulans*, a bacteria that has been implicated in sepsis among immunocompromised patients^54^. Remarkably, this bacteria was identified in 98% of ovarian RNA-Seq samples and represented by 555 unique peptides derived from 518 unique *N. circulans* genes (**Figure 6C**). In addition to its prevalence in ovarian cancer, 14 of these peptides were also consistently detected in esophageal cancer. As immunopeptidomic searches are known to inherently yield false positives, we searched the *N. circulans* proteome against eight neuroblastoma cell lines in which *N. circulans* RNA reads were not detected. None of the 14 peptides were found, suggesting that *N.circulans* was correctly identified at both the RNA and immunopeptidomic level across ovarian tumors (**Figure 6D**). We further validated these bacteria-derived peptides by comparing their MS/MS spectra to that of synthetic peptides, supporting the presence of these bacteria peptides in tumors (**Figure S9**). While we are unable to rule out the possibility that *N. circulans* is not a resident pathogen rather than a contaminant introduced during sequencing, the evidence at both the RNA and peptide levels across multiple datasets generated at different sites supports further investigation into the potential of targeting these pathogen-derived peptides, as well as the role of this bacterial strains in tumorigenesis. Collectively, we have systematically explored the landscape of pathogen-derived antigens in primary tumors and validated several novel antigens originating from oncogenic viruses or bacteria, highlighting the novel therapeutic potential of these pathogens when presented as tumor HLA peptides and introducing new avenues for investigation.

### Long interspersed nuclear element-1 (LINE1) ORF2-derived peptides are detected in immunopeptidome and represent pan-cancer targets

Transposable elements (TEs) are mobile genetic elements and constitute approximately half of the human genome. Recent studies have identified them as a novel class of pHLAs^55,56^. In healthy somatic tissues, TEs are tightly regulated epigenetically and their expression is typically silenced. However, this regulation is often lost in cancer leading to aberrant activation through epigenetic de-repression, positioning them as a novel source of tumor-specific antigens. TE expression and retrotransposition is generally associated with poor prognosis, contributing to genomic instability and accelerated tumor evolution^57^. More recently, immunopeptidomic research has focused on TE-derived chimeric transcripts, such as exonization and their ability to act as alternative promoter^58^, splicing donors and acceptors^56,59^ in order to expand their therapeutic potential. Notably, a recent pan-cancer analysis of TE-promoter events using RNA-Seq data identified 1,068 widespread tumor-specific aberrations, however, it remains unclear whether such events can generate detectable pHLAs in the primary tumor sample^55^. Moreover, certain TEs can encode proteins themselves, including Long Interspersed Nuclear Element (LINE), Long Terminal Repeat (LTR) elements such as endogenous retroviruses^60^, and sporadic reports of Alu, DNA transposons, and SINE-VNTR-Alus (SVA) retrotransposons^61^ (**Figure 7A,B**). We investigated the contribution of various TE categories to tumor-specific pHLA complexes and characterized their prevalence across cancers. Consistent with their ability to be translated from their cognate promoters, LINE1 and LTR elements generated the majority of self-translating TE-derived antigens, followed by DNA transposons. In contrast, SVA and SINE elements were more frequently involved in the formation of TE-chimeric transcripts (**Figure 7C, Table S9**).

**Figure 7.**
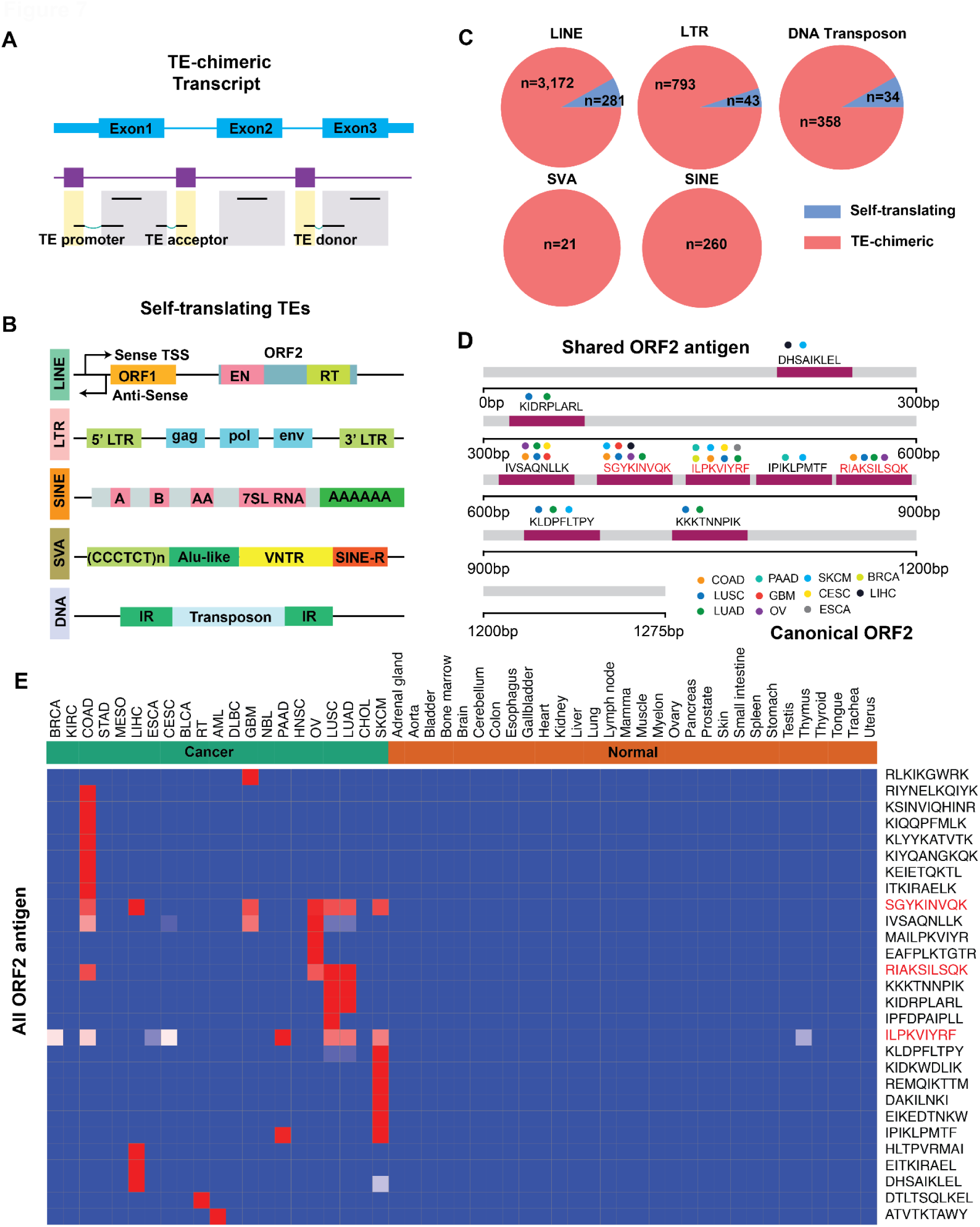
Landscape of Transposable Element (TE) derived antigens. (A,B) schematic overview of chimeric TE transcripts formed through exonization of de-repressed TE region in cancer (top), or potentially through self-translating mechanisms by utilizing their own promoter and coding regions (bottom). (C) number of TE antigens sourced from either self-translating TE or chimeric transcripts across five different classes of TEs (D) Detected pan-cancer peptide fragments from canonical ORF2 in immunopeptidome. (E) Tumor specificity of the ORF2 peptides across cancers and normal immunopeptidome. Validated ORF2 peptides are highlighted in red.

The ORF2 protein encoded by the LINE1 element plays a pivotal role in tumor evolution, functioning as both an endonuclease and reverse transcriptase to drive the retrotransposition of LINE1 and Alu elements. The endonuclease activity of ORF2 creates genomics instability, induces DNA double strand breaks and leads to large-scale genomic rearrangements and copy number alterations^62^. While LINE1 ORF2 protein has evaded detection through conventional methods such as western blotting and whole cell proteomics^63^, we surprisingly detected canonical ORF2 peptides recurrently presented on HLA across up to eight cancer types. The highest frequency of detection in colon, head and neck, and lung cancers (**Figure 7D, Table S9**). We confirmed the presentation of ORF2 peptides by synthesizing and validating three peptides recurrently identified from the immunopeptidome and demonstrated their tumor specificities (**Figure S9**, **Figure 7E**). To our knowledge, this is the first time that evidence of native ORF2 protein has been detected from endogenous expression across multiple cancers. This evidence suggests that ORF2 may be a short-lived protein that is rapidly degraded by the proteasome and presented on HLA before the full-length protein is detectable. Altogether, we assessed the potential of transposable elements to generate tumor-specific pHLAs through various mechanisms, and highlighted a previously elusive ORF2 protein, suggesting that immunopeptidomic characterization may represent an alternative approach for identifying difficult-to-detect proteins. These findings warrant further investigation into ORF2 and other TE biology, along with their utility as therapeutic targets.

## Discussion

Here we present the most comprehensive map of tumor-specific T cell targets to date, unveiling an abundant repertoire of previously uncharacterized derived from the “dark matter” of the immunopeptidome, representing actionable targets across the majority of human cancers. This work addresses a key bottleneck in immunotherapy development, the identification of tumor-specific antigens, by identifying potential pHLA targets in over 89% of cancers that substantially expands the cancer patient population that can benefit from T-cell-based therapies.

We mapped the cancer immunopeptidomic landscape using ImmunoVerse, an integrated computational pipeline capable of profiling 11 classes of molecular aberrations from transcriptomic data. Using this tool, we generated a catalog of non-canonical tumor-specific alterations by leveraging large-scale, histology-matched cancer RNA-Seq data, enabling us to construct a detailed and exhaustive search space for potential pHLAs for any immunopeptidomic sample. Integration of this pipeline with the Tesorai AI-based spectral search yielded a rich set of 28,446 tumor-specific pHLAs. These targets include those that span numerous tumor types, creating the potential for therapeutic development addressing large patient cohorts, shared mutation-derived neoantigens that can be rapidly incorporated in vaccine cocktails, and newly discovered targets that warrant further investigation.

To aid in the prioritization of therapeutic targets, we propose rigorous standards for defining actionable antigens in a therapeutic context, addressing this unresolved challenge by establishing a robust framework based on seven overarching criteria: (1) peptide abundance, (2) peptide recurrence, (3) absence in normal tissue RNA and immunopeptidome, (4) HLA binding affinity, (5) dependency of parent genes in tumor progression, and (6) homogeneity of target expression as measured by single-cell coverage and (7) confidence of peptide identification. Although current technologies and data availability limit our ability to apply all seven criteria to certain non-canonical antigen classes, our schema provides a roadmap for future investigations, helping tailor analyses to specific antigen types.

The search spaces we generated facilitate the integration of users’ own immunopeptidomic datasets, allowing a search across all classes of antigens even in the absence of matched RNA-seq data. Conversely, our tool is able to identify empirically supported pHLA candidates from RNA-seq data alone by referencing tumor-specific events catalogued in our atlas, enabling comprehensive characterization of high-confidence tumor targets from any RNA-seq or immunopeptidomic dataset.

Applying our analysis at a pan-cancer scale revealed numerous promising therapeutic targets. We identified 62 additional tumor-specific membrane proteins with favorable therapeutic windows for conventional CAR T cell targeting. Amongst the 28,446 tumor-specific pHLAs, we highlighted a highly recurrent, abundant, and tumor-specific HAVCR1-derived peptide in clear cell renal cell carcinoma, an ideal PC-CAR target, providing an alternative strategy to avoid the ectodomain shedding for conventional CAR.

We also report a spliced variant of the well-characterized melanoma-specific protein PMEL (sPMEL), which exhibits superior antigen abundance, tumor specificity, and recurrence across patients. Importantly, we demonstrate that sPMEL-specific T cells can be expanded *ex vivo*, highlighting its translational potential across multiple modalities including TCR-T, PC-CAR therapies, TILs expansion, bispecific T cell engagers and cancer vaccine development. We suggest that targeting sPMEL can lead to therapies that are safer, more potent and target a broader patient population than current clinical PMEL-directed immunotherapies, illustrating the utility of our target prioritization framework for refining antigen selection even among highly similar tumor targets.

In addition, we discovered dozens of pan-cancer cryptic-ORF derived tumor-specific antigens amenable for PC-CAR targeting. These targets may be particularly important targets in the immunologically cold tumors where conventional neoantigens have been insufficient to achieve therapeutic efficacy.

Our immunopeptidome-guided search for mutation-derived neoantigens reveals numerous novel neoantigens in colon cancer and liver cancers, several of which we have validated experimentally. Unlike prediction-based approaches that yield high false-positive rates, our analysis provides empirical evidence of peptide presentation for these neoantigens, supporting their inclusion in the current vaccine cocktail. To enable rapid and high-confidence neoantigen selection, we have incorporated a feature into ImmunoVerse that allows the identification of empirically “presentable” neoantigens based on RNA-seq data alone, which we expect to improve the prioritization of naturally presented peptides that are likely to induce immunogenicity. We anticipate that this approach will enhance the precision and clinical efficacy of personalized cancer vaccines In addition, we revealed a rich set of peptides derived from tumor-resident pathogens, including known oncogenic viruses and bacteria and newly identified pathogens. We highlighted recurrent previously unreported CMV-derived peptides presented across tumors as well as new peptides derived from known tumorigenic pathogens such as *F. Nucleatum* that can provide potential therapeutic avenues against tumors. Strikingly, we detected an abundance of peptides derived from the underexplored bacterium *N. circulans* detected in 98% of ovarian cancer patients, warranting additional validation and investigation into the biological role of this species in ovarian cancer initiation. Together, these pathogen-derived peptides represent a previously underappreciated source of new high-confidence targets for new candidates for therapeutic exploitation.

Our analysis also yields novel biological insights into the HLA presentation pathway and supports the preferential presentation of defective and short-lived peptides such as splicing products and cryptic ORFs. These findings, together with our identification of previously elusive ORF2 peptides from the LINE1 element, are consistent with the hypothesis that Defective Ribosomal Products (DRiPs) may be overrepresented in HLA-presented peptide repertoire^64^, highlighting the diverse and dynamic nature of antigen presentation and suggesting that immunopeptidomics could be used as an alternative strategy to identify difficult-to-detect unstable proteins. While we highlighted several biological insights that are gleaned from these resources, we expect that additional analyses of these datasets can inform numerous new avenues of investigation.

While our analysis focused on identifying intrinsically presented tumor antigens by HLA class I-restricted tumor antigens, our tools can also be adapted into other settings. Our pipeline can be applied to identify HLA-II antigens on tumor surfaces for immune surveillance, which are expressed in many cancer types and remain underexplored^65^. Lastly, our pipeline is highly adaptable, allowing the integration of additional modalities such as whole-genome sequencing (WGS) and Ribo-Seq, as well as new molecular classes, including circular RNAs and RNA editing.

We acknowledge several limitations in the current study. We highlight that the availability of immunopeptidome data varies across cancer types; for instance, in bladder cancer, certain tumor-specific genes like VGLL1 are not identified in the current immunopeptidomic data, potentially due to limited tumor samples and HLA alleles represented in publicly available samples. We anticipate that expanding the pool of immunopeptidomic datasets in future studies will help bridge existing knowledge gaps. Moreover, while our analysis has delineated a cancer-specific search space, a more granular examination of individual tumor subtypes may uncover additional vulnerabilities that are not captured in our broader analysis. Additionally, as our atlas is primarily based on bulk sequencing data, these samples may include signals from contaminating stromal or immune cells in both normal and cancerous tissues, and do not capture intra-tumoral heterogeneity. As such, the results presented in the portal should be interpreted within the appropriate biological context.

Additional classes of alterations including those deriving from proteasomal splicing^66^, post-translational modifications (PTMs)^17^ and alternative codon^67^ can generate pHLAs as well. The challenges for targeting these are two-fold. First, since these events can arise in any place and in random combinations, enumerating all possibilities has been computationally prohibitive. Second, defining tumor specificity of post-translational events is non-trivial, as these events are poorly cataloged in normal tissues and may also occur under various stress conditions. Future research dedicated to rigorously evaluating the therapeutic value of these novel classes of antigens could further significantly expand the repertoire of tumor antigens.

From this rich dataset, we reveal actionable targets in 89% of tumors and highlight several examples of promising therapeutic avenues, including membrane proteins, differentially expressed pHLAs, tumor-specific splicing events, TEs and through peptides derived from tumor-specific pathogens. We validated a panel of 19 targets across six antigen classes and demonstrated anti-tumor T cell activity through these targets, underscoring the high confidence of peptides identified through our pipeline and the immediate therapeutic potential of these findings. By making the ImmunoVerse platform freely available both as a web portal and cloud-based pipeline, we democratize access to immunotherapy target discovery for the global research community. We envision that these findings and resources will provide a foundational roadmap for prioritizing the next generation of immunotherapy targets, unlocking the development of novel immunotherapies that offer safe and effective new options to a broad range of cancer patients.

## Acknowledgements

We would like to thank Drs. Alvin Farrel and John M. Maris for providing the neuroblastoma PDX Ribo-Seq data. We also thank Drs. Rowan Beck and Cera Fisher from Velsera to provide technical support for running the pipeline on the Cancer Genome Cloud (CGC) platform. We thank Drs. Shohei Koide, Akiko Koide and Lonrezo Maso and Epsa Rajak for sharing the HLA and β2M constructs and for assistance in pMHC preparation. We thank Dr. Ritchlynn Aranha for the valuable inputs on the HLA presentation pathway. We thank Dr. Marcin Imieliński for the assistance with the neoantigen cancer vaccine analysis. The mass spectrometry experiment is supported in part by NYU Langone Health and the Laura and Isaac Perlmutter Cancer Center support grant P30CA016087 from the National Cancer Institute experiment. This work was delivered as part of the NexTGen team supported by the Cancer Grand Challenges partnership funded by Cancer Research UK (CGCATF-2021/100038 to M.Y) and the National Cancer Institute (CA290738-01 to M.Y.) and The Mark Foundation for Cancer Research, NIH Director’s New Innovator Award (1DP2CA301080 to M.Y.), Cancer Genome Cloud (CGC) collaborative fund (G.L and M.Y.). M.Y. is the Bakewell Foundation-Rachleff Innovator of the Damon Runyon Cancer Research Foundation (DRR-86-25).

## Data Availability

The curated molecular catalogs of tumor-specific aberrations, scripts used to generate the figures and tumor-specific search space are available at https://github.com/frankligy/pan_cancer_intracellular_antigen_atlas, and the pipeline for generating the customized search space is hosted on the Cancer Genome Cloud with tutorials (https://frankligy.github.io/pan_cancer_intracellular_antigen_atlas/notes/deployed_pipeline_latest.pdf). Reviewers can access the CGC platform (https://cgc.sbgenomics.com/home) using following credentials (username: reviewer, password: ImmunoVerse3). The ImmunoVerse web portal is accessible through (https://www.immuno-verse.com/) or alternatively through (https://immunoverse-web-5ef5cdbdcedf.herokuapp.com/).

## Contributions

G.L. and M.Y. conceived and designed the study. G.L. developed the computational pipeline, constructed the atlas, and conducted all bioinformatics analyses. O.U.G. carried out peptide spike-in validation and refolding experiments. A.S. assisted in analyzing the normal immunopeptidome data and refolding experiments. Z.H. performed single-cell analysis for self-antigens. X.L. contributed to modeling the protein structure. D.S, M.B. P.C. analyzed immunopeptidome data through Tesorai Search platform. I.C. and B.U. performed synthetic peptide validation experiments and validation. D.R. analyzed the sequencing data from the Hartwig Medical Foundation cohort. G.L, M.D., R.C.C and M.Y. designed the experiments. E.T., A.N., and M.K. performed the PMEL TCR cross-reactivity assays. D.T. provided critical insights into the analysis of Transposable Elements. P.S., R.Y., M.D., R.C.C. executed the PMEL validation experiments. G.L. and M.Y. wrote the manuscript.

## Conflict of Interests

M.Y. is a co-founder and holds equity in HuLA Therapeutics. D.S., M.B., and P.C. are salaried employees of Tesorai and hold stock in the company. G.L and M.Y. have filed PCT patents for the bioinformatics pipeline and the targets discovered in the paper.

## Method

### Collection of public bulk RNA-Seq

The tumor bulk RNA-Seq data was collected from TCGA (phs000178.v11) and TARGET (phs000218.v26) upon dbGaP approval through Cancer Genome Cloud (CGC) platform which deposited uniformly processed BAM files aligned to Hg38 human genome assembly using STAR 2-pass mode. 21 adult and pediatric tumor types were selected based on the availability of matched immunopeptidome public datasets. The normal tissue bulk RNA-Seq raw data was downloaded from NCBI SRA upon dbGap approval (phs000424.v6.p1). The normal tissue gene expression data (GTEx_Analysis_2017-06-05_v8_RNASeQCv1.1.9_gene_tpm.gct.gz) was retrieved from the GTEx portal (https://gtexportal.org/home/downloads/adult-gtex/bulk_tissue_expression) as pre-calculated Transcripts Per Millions (TPMs) values.

### Collection of public immunopeptidome data

Publicly available immunopeptidome data sets were manually curated and downloaded from both PRIDE (https://www.ebi.ac.uk/pride/) and Massive platform (https://massive.ucsd.edu/ProteoSAFe/static/massive.jsp). The biological context associated with each immunopeptidome experiment was retrieved from the original publications, along with the reported HLA alleles wherever possible. Normal tissue immunopeptidome raw data, sample annotations and HLA alleles were downloaded from HLA ligand atlas with the accession number PXD019643.

### Collection of public tumor single-cell RNA-Seq data

Publically available tumor single-cell RNA-Seq for 14 cancer types (thyroid carcinoma, colon cancer, gastric cancer, bladder cancer, head and neck cancer, pancreatic adenocarcinoma, ovarian cancer, lung squamous cell carcinoma, lung adenocarcinoma, glioblastoma, sarcoma, rhabdoid tumor, Uterine corpus endometrial carcinoma, breast cancer) were downloaded from Cancer Single-Cell Expression map (CancerSCEM v1.0)^68^ (https://ngdc.cncb.ac.cn/cancerscem/). Uniformly processed h5ad files were downloaded and the malignant cells were selected based on marker genes expression (EPCAM, FOLH1, KLK3, KRT8, KRT18, KRT19) as documented by CancerSCEM official tutorial. The average gene expression across malignant cells was calculated, ranked (in ascending order) and the percentile of each gene across all detected genes were reported as the single-cell coverage (homogeneity) in the analysis.

### Computational Pipeline to identify diverse tumor-specific molecular events

A comprehensive computational pipeline was developed to profile 11 classes of molecular aberrations in the tumor, filtered by normal tissue presence, and assembled into sample-specific or tumor-specific peptide search space for downstream immunopeptidome analysis. The detailed descriptions of each module were as following:

1. Self-gene: For any user-supplied data, the raw RNA-Seq fastq files were subject to Kallisto (version 0.44.0) pipeline^69^ using transcriptome index generated from the human Gencode V36. Transcripts Per Millions (TPMs) for each gene were calculated by summing up all transcript-level TPMs associated with each gene. The normal tissue safety profiles for each gene were retrieved from BayesTS database (https://github.com/frankligy/BayesTS/blob/main/database/full_results_XY_essential_tiss <u>ues.txt</u>) based on multiple evidence including normal tissue gene expression, protein staining and essential tissue distributions^3^. The median TPM in each normal tissue is calculated, and the maximum of them serves as a proxy for normal tissue expression for filtering purposes. Next, A gene is defined as tumor specific only if met following criteria: (a) median TPM in tumor is greater than 20, (b) median TPM in tumor should be greater than the normal expression calculated as above, (c) BayesTS score is less than 0.3 to ensure safety of targeting it therapeutically, (d) significantly upregulated in the tumor versus histology-matched normal tissue when the differential gene analysis data is available. Identical selection criteria were applied to the selection of surface protein targets. While these selection criteria effectively filtered out the majority of non–tumor-specific genes, we conducted post hoc manual inspection of all retained targets and further excluded those with appreciable expression in essential normal tissues (blacklist genes; **Table S5**). For the TCGA and TARGET data, we retrieved the gene expression TPMs from Xena Browser (https://xenabrowser.net/datapages) for each cancer, the differentially gene expression between tumor versus histology-matched controls were retrieved from GEPIA^70^ API (http://gepia.cancer-pku.cn/detail.php), and the genes with adjusted p-value less than 0.05 and log fold change (LFC) greater than 0.58 were defined as significantly upregulated genes in each tumor. The protein sequence of canonical self-genes and isoforms were downloaded from Ensembl and Uniprot, respectively. We note slight differences between ensembl-annotated main transcript and Uniprot-annotated main transcript for a subset of genes, thus utilizing the references from both sources can cover all the reviewed protein coding isoforms. Using the tumor gene expression, we filtered out proteins and isoforms that are not expressed in the tumor sample., and only the expressed proteins and isoforms were assembled as either sample or tumor-specific search space in fasta format.
2. Alternative Splicing: For user-supplied RNA-Seq raw data, the fastq files were first aligned to human hg38 reference using STAR 2-pass mode to generate the BAM file^71^. pysam package (https://github.com/pysam-developers/pysam) was utilized to identify all the junction reads (spanning the introns based on the N in the CIGAR string) and quantify the read counts that support each splicing junction (find_introns function). The comprehensive GTEx normal tissue splicing junctions profiles were downloaded from SNAF database (http://altanalyze.org/SNAF/download.tar.gz)^14^. A splicing junction was defined as tumor-specific only if met following criteria: (a) average read count in tumor is greater than 10, (b) average read count in all normal tissues is less than 1, (c) the log fold change between tumor and normal tissues is greater than 4. To further remove potential alignment error from genuine splicing events, splicing junctions with no annotated splicing sites will be discarded. For TCGA and TARGET data, pysam packages were applied on the CGC uniformed processed BAM files from STAR 2-pass mode. For the pan-cancer analysis, since the RNA-Seq and immunopeptidome are not matched by exact sample but only by histology, frequently detected splicing junctions (frequency > 20%) were retained as they should be considered as shared events in each tumor type instead of specific sample. The tumor-specific splicing junctions and the flanking sequences (n=33nt) were further subject to *in silico* translations, potential splicing peptides were defined as ones without stop codon. When the tumor-specific splicing junction coincides with annotated isoforms in Uniprot, the whole documented isoform sequences were used instead. All the tumor-specific isoforms and tumor-specific local junction derided peptides were assembled into sample-specific or tumor-specific search space.
3. Intron Retention: The intron regions were defined based on previously described exon-intron model^14^ by first collapsing all the annotated isoforms in Ensembl associated with each gene, only regions that are consistently annotated as intron across all isoforms will be considered. Introns that are overlapping with exons from another gene were also discarded because the reads coverage likely come from the exons not the introns. Next, Stringtie2 (version 2.1.3) was run in de novo mode^72^ to predict all the potential novel transcripts including the ones with intron retentions. When a defined intron was supported by a predicted read-through Stringtie transcript, it was considered as an intron-retention candidate, and the expression (TPM) of the exon covering the intron regions was utilized as a proxy for the expression of that intron retention event. Exact same pipeline was applied to the GTEx normal samples across 31 histologies. We randomly subsample 5 normal samples from each histology as the representative intron retention profiles in each tissue. Only intron retentions that are absolutely absent in normal tissue were retained as tumor-specific intron intentions. We further utilized the previously reported SNAF database for intron retention detected from GTEx to make sure all the intron retention considered are not present in the normal tissues. For pan-cancer analysis, same as the splicing event, we defined frequently identified intron retention events (frequency > 15%) in the tumor RNA-Seq cohort to include in the immunopeptidome search. Next, we utilized ORFfinder program downloaded from NCBI (https://www.ncbi.nlm.nih.gov/orffinder/) in default setting to predict all the potential ORFs that can be generated from the intron-retained transcript, and the most probable ORF was ranked primarily by the length of the ORF. When the length difference between the best-scored ORF and the second-best scored ORFs is less than 8nt, we further ranked them based on the GC content and the codon potential in humans, as described before^14^. The most probable ORF was identified and translated into peptide. Lastly, only peptides that overlap with the intron region of interest were considered as the causal peptides originating from the intron and were reported in the sample-specific or tumor-specific search space.
4. Tumor-resident microbiome: The unmapped reads in the tumor BAM files were collected using samtools^73^ and reverted back to the fastq format using bedtools^74^. The fastq files were then subject to the kraken2 program^75^ to be classified into hierarchical taxonomies, using a comprehensive indexed database (PlusPF) downloaded from (https://benlangmead.github.io/aws-indexes/k2), which was constructed using archaea, bacteria, virus, plasmid, human univec_core, protozoa and fungi reference sequences. Importantly, the human genome was also included to further remove any remaining human reads. While there is no consensus cutoff for kranken2 program and is heavily dependent on the specific use case, to mitigate the false discoveries rate, we adopted a stringent cutoff (confidence=0.9) such that a read would only be assigned to taxonomy if over 90% of the consecutive kmers can be unambiguously mapped to it. Similar to the intron retention, a representative normal cohort was compiled and the exact same pipeline was applied to remove pathogens detected in normal tissue. Tumor-specific microbiomes were defined as species that are not detected at species level in normal control. To increase stringency, species belonging to genera with appreciable levels of mapped reads were also filtered out. For the TCGA data, the frequently detected microbiomes (frequency > 20%) were selected, and were further cross-referenced with the published TCMbio atlas^50^ that utilized the same datasets with dedicated quality control steps to remove potential contaminating reads and a bayesian inference step to distribute ambiguous reads mapped to parental taxonomies, potentially leading to more accurate quantifications^76^. Microbiomes that are recurrently detected in both two procedures were selected for histology-matched immunopeptidome search. The reference proteomes for the selected microbiome were retrieved from Uniprot, when multiple reference proteomes were available, we compared the completeness of the proteomes and selected the most complete one. The microbiome search space was further appended by the literature-supported tumor-resident virus, or when the immunopeptidome datasets were known to harbor additional microbiomes.
5. Transposable element (TE)-chimeric transcript: The exonization of TE into a spliced transcripts can manifest through either TE-promoter, or TE splicing site donor/acceptor. Firstly, the 8 prebuilt TE locus index for human hg39 reference were downloaded from TElocus^77^ database (https://labshare.cshl.edu/shares/mhammelllab/www-data/TElocal/prebuilt_indices), along with the known genes from hg38 downloaded from UCSC (https://hgdownload.soe.ucsc.edu/goldenPath/hg38/bigZips/genes/) and the chromosome coordinate annotation table (https://labshare.cshl.edu/shares/mhammelllab/www-data/TElocal/annotation_tables/). Following the previously reported procedures^55,56^, the tumor-specific splicing junctions defined above were further annotated based on whether the splicing sites fall into annotated TE regions, and the TE regions following the splicing site till the end were included in the spliced transcript. Both sense and antisense strands of each TE region were considered for TE-chimeric transcripts. Similar to the *in silico* translation protocol described for local splicing junctions, TE-chimeric regions including both the canonical exons and the TE regions were translated and assembled into sample-specific or tumor-specific search space.
6. Autonomous transposable element (TE): Similar to TE-chimeric transcript pipeline, annotated TE regions were quantified by TElocus program. Due to the nature of highly repetitive sequences associated with TE, TElocus efficiently utilized both uniquely mapped and multi-mapped reads to re-distribute the RNA-Seq reads previously mapped to the gene-only reference. The TE-locus reported raw counts were further normalized by sample-specific sequencing depth into Counts Per Millions (CPMs), as described before^78^. Similar to intron retention and tumor-resident microbiome analysis, normal tissue cohorts were assembled to help filter out non-tumor-specific TEs. An autonomous TE was defined as tumor specific when the median CPM in tumor is greater than 1 and the mean CPM in normal tissue is less than 0.5, together with the log fold change greater than 5. These cutoffs were selected based on empirical observation between tumor and normal TE expressions, and the resultant TE events following this protocol often possess desirable therapeutic windows. For TCGA and TARGET analysis, detection over 20% of tumor patients from RNA-Seq were added for the immunopeptidome search. Amongst all the TE classes and TE families, the Long Terminal Repeats (LTRs) including Human Endogenous Retrovirus (HERVs), Long Interspersed Nuclear Element (LINE), Short Interspersed Nuclear Element (SINE), DNA transposon and SVA family from retroposon class were selected as potential translatable TEs. Both sense and antisense directions were considered for *in silico* translation.
7. Gene fusion: The user-supplied RNA-Seq fastq files were analyzed by STARfusion^79^ (version 1.13.0) packaged as a docker container downloadable from (https://data.broadinstitute.org/Trinity/CTAT_SINGULARITY/STAR-Fusion/). The STARfusion database was downloaded from (https://data.broadinstitute.org/Trinity/CTAT_RESOURCE_LIB/GRCh38_gencode_v44_C <u>TAT_lib_Oct292023.plug-n-play.tar.gz</u>), and was executed in the default setting documented on the official tutorials. The reported in-frame fusion transcripts that are not labelled as (1) GTEx_recurrent_StarF2019, (2) BodyMap, (3) DGD_PARALOGS, (4) HGNC_GENEFAM, (5) Greger_Normal, (6) Babiceanu_Normal, (7) ConjoinG were considered tumor-specific fusion events, and the fusion peptides were used as the sample or tumor-specific search space. For TCGA data, the previously curated gene fusions were retrieved from ChimerDB (https://www.kobic.re.kr/chimerdb/) with both detected fusions from different softwares^80^. The chromosomal coordinates of each gene fusion were extracted from the downloaded fusion result files and were further translated to fusion peptides. Fusions detected in TCGA matched paratumor samples were filtered out due to their suspicious tumor-specificities. For TARGET data, the CGC uniformly processed gene fusion files from STARfusion were downloaded. Following the same procedures, chromosomal coordinates of the fusion breakpoints were utilized to retrieve the fusion flanking nucleotides sequence and to derive the fusion junction peptides.
8. Variants and somatic mutation: For user-supplied RNA-Seq raw data, one of the recommended RNA variant detection protocols that achieves superior performance in the benchmark were used to identify variants from RNA-Seq reads^81^. Specifically, Opossum^81^ was applied to the raw BAM files to conduct a series of quality controls, rendering the resultant BAM files suitable for variant caller. It includes (a) removing low quality and improperly-aligned reads, (b) deduplication, (c) collapsing overlapping reads, (d) separating intron-spanning reads into multiple segments. All the parameters were left as default except the MinFlankStart and MinFlankEnd were set as 10 to remove spurious bases at the start or end of aligning reads, a common artifact in calling variants from genomics and transcriptomics data^82^. Subsequently, the modified BAM files were analyzed by Platypus^83^, a haplotype-based variant caller to report high-confidence variants in the tumor samples. Variant Effect Predictor (VEP) was applied to characterize the impact of each variant on protein coding regions^84^. The missense variant, inframe deletion and insertion, along with frameshift variants were retained. While definitely calling somatic mutations from tumor samples alone without the sample-matched normal control is impossible, certain predictors have been proposed that can achieve reasonable prediction power to distinguish somatic mutation from germline polymorphisms^85^. A variant was predicted as likely somatic mutation if met all following criteria: (a) absence or extremely low population frequency (<0.0001) in dbSNP, (b) absence in normal tissue RNA editing database REDIportal^86^, (c) maximum variant allele frequencies in tumor samples less than 0.95. When a variant is confirmed as somatic mutation from the external COSMIC database, it was retained as well^87^. While these variants were annotated as likely somatic mutations, all the variants will be utilized to construct the search space as germline polymorphism also directly affects the protein sequence and should be considered. Additionally, all the detected A to G variants were annotated as likely RNA editing events in the final results. For TCGA data, somatic mutations called from Whole Exome/Genome Sequencing with matched normal samples were downloaded from Xena Browser (https://xenabrowser.net/datapages/). For TARGET data, the uniformly processed somatic mutation vcf files were downloaded from CGC platforms. Due to lack of mutation data in TARGET for rhabdoid tumor, a recently published rhabdoid tumor WGS study^88^ was used instead to impute the somatic mutation burden for this tumor type.
9. lncRNA, (10) Pseudogene and (11) cryptic open-reading frames (ORFs): Since RNA-Seq usually lack the capacity to comprehensively quantify the underlying lncRNA and pseudogene expression levels, due to suboptimal capture protocol. A Ribo-Seq informed database nuORF^16^ (version 1.0) was used as a more comprehensive catalogue for likely translatable lncRNA, pseudogene and unannotated open-reading frames either residing in the 3’ /5’ untranslated regions or overlapping with annotated ORF but adopting a different reading frames. Using the provided hg19 transcripts and chromosomal coordinates information in the original nuORF database, we first rederived all the ORF peptide sequences and assured the correct length compared to the original paper. These nuORF peptide databases were appended to all the immunopeptidome searches to enable the identification of these three molecular sources.

### Immunopeptidome data Analysis

The exhaustive tumor-specific search space was generated by incorporating all tumor-specific molecular aberrations to search the histology-matched large-scale immunopeptidome datasets using MaxQuant^19^ (version 2.4.9.0). MaxQuant was run in nonspecific enzyme mode with peptide length ranging from 8-11 mer, a typical setting for HLA-I peptide discovery described in numerous previous studies^89–91^. Match between run and de novo sequencing were enabled for each search by default. Particularly, no peptide FDR was enforced in the initial run to obtain the full list of peptide-spectrum matches (PSMs). The full PSM list was then subject to MS2Rescore progrem^20^, which has been reported to increase the immunopeptidome identification rate by 46% using additional features and deep-learning predicted spectral libraries, to rescore the original p-value and re-rank the PSM lists. Subsequently, a 5% peptide-level FDR, typically used in immunopeptidome search and has been shown to rescue bona fide antigens which would otherwise be missed at 1% FDR^92,93^ was applied to both MaxQuant reported Posterior Error Probability (PEP) and MS2Rescore reported q-value to get the FDR-controlled peptide lists. Protein-level FDR was not applied, as the primary focus of this analysis is on HLA-presented peptides. We used similar search parameters with Tesorai Search, with the exception of using 1% peptide-level FDR and further included longer HLA-I peptides (8-15mer). The union of identifications from both search engines was summarized for each tumor immunopeptidome dataset to generate the initial list of evidence-supported peptides. In cases where a single MS/MS spectrum matched different peptides in MaxQuant and Tesorai Search, we applied the following criteria to resolve conflicts: (1) If the Andromeda score was >70 and the Tesorai score <5, the MaxQuant identification was retained; (2) if the Andromeda score was <70 and the Tesorai score >5, the Tesorai identification was retained; (3) if both scores were high (Andromeda >70 and Tesorai >5), the identification was considered ambiguous and excluded; (4) if both scores were low (Andromeda <70 and Tesorai <5), the identification was deemed low confidence and also excluded. The cutoffs for both Andromeda (>70) and Tesorai Search (>5) were determined in consultation with the original developers, supported by extensive manual inspection of MS/MS spectra by both teams to reach consensus on the selection criteria.

Next, netMHCpan 4.1^22^ was employed to predict the binding affinity between each identified peptide and the cognate HLA alleles from the sample it was identified, both strong (rank < 0.5%) and weak binders (rank < 2%) were considered, and only the peptides predicted to bind with cognate HLAs were retained to further remove possible false discovery. The maxquant-reported precursor intensity associated with each PSM was reported to estimate the peptide abundance in each immunopeptidome experiment, when multiple scans mapped to the same peptide, maximum precursor intensity was used as the peptide abundance estimate due to its high correlation with the area under the curve (AUC) reported by proprietary software (**Figure S10**). When comparing peptide abundance across different peptides, quantile normalization was conducted on the raw intensity values. Briefly, all the eluted peptides in each immunopeptidome experiment were ranked in descending order based on precursor intensity values, the upper quantile (75%) value as selected as the base, and the raw intensity of each peptide was divided by the base value and then log2-converted. For safety screening of the identified tumor-specific antigens, normal immunopeptidome raw data was obtained from the HLA Ligand Atlas (PXD019643) and reanalyzed using both MaxQuant and Tesorai Search with search parameters identical to those used for tumor samples. Peptides detected in any normal tissue were annotated, and those identified in essential tissues (excluding adrenal gland, ovary, prostate, testis, and thymus) were excluded from the final list due to potential off-tumor toxicities.The absence of Cryptic-ORF-derived peptides was further cross-referenced to IEAtlas database^94^ to ensure tumor specificity.

### Alphafold2 modeling

The non-docker version of Alphafold2^95^ (version 2.3.0) was downloaded and installed from (https://github.com/kalininalab/alphafold_non_docker). The HLA alleles sequences were retrieved from IMGT/HLA database (https://www.ebi.ac.uk/ipd/imgt/hla/). Signal peptides predicted by SignalP^96^ (version 5.0) were removed from the N-terminus and the remaining HLA peptide sequence was served as input, together with β2M and peptide sequence. Alphafold2 was run in multimeter mode without relaxation, and the -t argument was wet as 2023-04-29. The highest ranked PDB files (rank_0.pdb) were used for visualization in Pymol3 software (http://www.pymol.org/pymol).

### Ribo-Seq Analysis

Ribo-seq for 8 neuroblastoma PDX models was conducted by the Maris lab at Children’s Hospital of Philadelphia (CHOP). The Ribo-Seq raw fastq files were analyzed by the RibORF (version 2.0) pipeline^97^. Briefly, the adapter sequence (TGGAATTCTCGGGTGCCAAGG) was removed from the pair-end Ribo-Seq files. Bowtie2^98^ (version 2.3.1) was used to first build the ribosomal RNA index and to align the raw reads to the rRNA reference, removing the noninformative reads. Reads that cannot be mapped to the rRNA reference were subsequently aligned to the human hg19 genome by TopHat^99^ to get the SAM file. For each Ribo-Seq read length, a diagnostic plot was automatically generated by the RibORF pipeline for users to specify the optimal offset length to correct the ribosome length, which is normally 12 in the human. The specified offsets were further used as input to convert each aligned ribosomal read to its A site to obtain the simplified SAM file. Both the simplified and original SAM files were directly visualized in Integrative Genomics Viewer (IGV) to evaluate the ribosomal occupancy of selected high confidence cryptic ORF regions. Further, the translatable probability of each cryptic ORFs was reported by RibORF. Both the visual inspection and the quantitative translatable likelihood were considered to decide whether a region is considered translatable in neuroblastoma PDX models.

### Clustering and visualization of RNA and immunopeptidome datasets

To cluster all public RNA-Seq samples from the collected TCGA and TARGET cohorts, transcript-per-million (TPM) values of 13,033 genes in which each can give rise to at least one uniquely self peptide detected in our atlas were used as input features. For clustering public immunopeptidome datasets, all eluted peptides, including both tumor-specific and non-tumor-specific peptides derived from canonical protein-coding genes, were included. The feature matrix was constructed using the median peptide abundance for each parental gene when multiple peptides from one gene are identified, the peptide abundance is expressed as descending percentiles ranging from 0 to 1, where 1 indicates the most abundant peptide within a sample and 0 the least. For both RNA and immunopeptidome clustering, the Scanpy^100^ package was used for downstream analysis with following steps and parameters: the top 5,000 highly variable features were selected using the highly_variable_genes function with the “seurat” method, followed by scaling (with max_value=10), principal component analysis (PCA, with 50 components), neighborhood graph computation using default parameters, and Uniform Manifold Approximation and Projection (UMAP) coordinate calculation also with default settings.

### *N. circulans* Alignment Validation

Three tumor RNA-Seq samples were randomly selected from ovarian cancer (OV), esophageal cancer (ESCA), and stomach cancer (STAD) in which *N. circulans* were evidenced. To avoid the detection purely due to potential batch effects, samples from different institutions inferred by their TCGA sample IDs were prioritized (**Table S8**). TCGA-provided BAM files were downloaded from the Cancer Genome Cloud (CGC) and re-analyzed using pathogen detection modules. The reference genome (FASTA) and annotation file (GTF) for the *N. circulans* reference strain submitted by US FDA were obtained from the NCBI website (https://www.ncbi.nlm.nih.gov/datasets/genome/GCF_013267435.1/). Only paired-end reads uniquely mapped to *N. circulans* by Kraken2 (Taxid=1397, confidence threshold >90%) and fully aligned to the reference genome were retained. The genomic coordinates of these reads were recorded and visualized using pyCircos (https://github.com/ponnhide/pyCircos). The 16S rRNA regions were annotated using the corresponding reference GTF files and visualized by pyCircos. On the protein level, due to discrepancies between the *N. circulans* proteome available in UniProt (used in the pan-cancer search) and the proteome of the US FDA reference strain in NCBI, a customized proteome was generated for the purpose of this alignment validation. Specifically, CDS features from the GTF file of the FDA strain were extracted and only coding sequences with lengths divisible by three were retained and translated *in silico* using the standard prokaryotic codon table (NCBI Codon Table 11). Only full-length protein sequences without premature stop codons were kept. This curated NCBI-derived proteome was then searched against all publicly available ovarian cancer immunopeptidomes using Tesorai search platform leveraging its high scalability and processing speed. Matched peptides and their genomic locations were visualized using pyCircos for ovarian cancer samples.

### Peptide validation

Twenty peptide candidates across different molecular classes were synthesized (GenScript, Piscataway, NJ) at the minimum of 95% purity with an average yield of 5-9 mg. Peptides were reconstituted with either 100% water, 5% DMSO/95% water or 0.1% HCOOH/99.9% water to a working concentration of 2mM and further diluted using HPLC-grade water to a final stock concentration of 10pmol/uL. 2 pmol of each peptide was analyzed using an Evosep One coupled to an Orbitrap Tribrid Eclipse (Thermo Fisher Scientific) using a 30 SPD gradient (Evosep; column EV1106). MS methods including mass analyzer, resolution, quadrupole isolation window, activation type, and activation energy were matched exactly to that of the original immunopeptidomic analysis, with the exception of orbitrap resolution 17.5k which was changed to 15k. For two peptides (QPKTKLLLL and KLKPGILKK; **Figure S9**) the Wideband Activation feature on LTQ-type Thermo Fisher instruments was used for the original tumor immunopeptidomic analysis. This feature is not available on the Eclipse Tribrid, so Multistage Activation was used instead, activating also the M-17 ion. MS/MS spectra of the synthetic peptides were visualized using Xcalibur Qual browser (Thermo Fisher Scientific) and the spectrum with the highest normalized level (NL) intensity was selected as the representative spectrum, and the matched spectra were visualized using spectrum_utils^101^ (https://spectrum-utils.readthedocs.io/en/latest/plotting.html) and pyteomics packages (https://github.com/levitsky/pyteomics). All matched spectra were manually validated to confirm the antigen identification.

### Peptide MHC (pMHC) refolding

The recombinant HLA heavy chain and beta-2-microglobulin (β2m) were produced using the pET28 and pHFT2 vectors, respectively, with their expression and purification carried out according to established protocols described in earlier studies^102,103^. HLA construct has a His6-tag and Avi-Tag (Avidity). HLA protein was expressed and in vivo Biotinylated using inclusion bodies in Escherichia coli BL21(DE3) co-expressing BirA enzyme and adding 50 µM of biotin in Luria Broth media. β2m construct contains His6-Tag and TEV protease site, this last feature is to remove the His-tag previous refolding, as HLA expression B2m was expressed using inclusion bodies in E. coli BL21(DE3). Inclusion bodies were solubilized by using Urea 8M buffer, Urea-soluble fraction was subjected to affinity chromatography using Econo-Pac Chromatography Columns (Bio-Rad, cat. no. 7321010) packed with Ni Sepharose 6 Fast Flow histidine-tagged protein purification resin (Cytiva, cat. no. 17531801); HLA bound to the resin was eluted with 10 mL of urea buffer containing 0.5 M imidazole. A diafiltration step with Urea buffer was performed to remove the Imidazole, and the protein was further concentrated to a final concentration of >100 µM.

β2m purification was performed using a similar protocol to that of HLA, with a key modification to include an on-column refolding step, which was achieved by passing a gradient of Urea buffer (8 M urea, 0.1 M Tris-HCl, pH 8) and Native buffer (50 mM Tris-HCl, 250 mM NaCl, pH 8) through the column. The gradient was applied to gradually reduce the urea concentration until only native buffer was used. β2m was eluted with Native buffer + 0.5 M Imidazole, to remove Imidazole and separate persistent aggregates from the monomeric fraction, the concentrated β2m solution was loaded onto the AKTA Pure system and subjected to size exclusion chromatography (SEC) using a Superdex 75 Increase 10/300 GL column (Cytiva, cat. no. 29148721). Monomeric β2m fractions were collected for further processing. His-tag on B2m was cleaved using TEV protease (Sigma, cat. no. T4455), and following cleavage, the protein mixture was run through the AKTA system one more time to isolate the cleaved β2m monomer. The purified cleaved monomeric β2m was then used for peptide-MHC refolding.

Synthetic peptides were dissolved to a final concentration of 2 mM in the appropriate buffer, as determined by a qualitative solubility test provided by GenScript. Refolding of the peptide-MHC (pMHC) complex was performed in the PBS buffer using 3 µM cleaved β2m, 30 µM peptide, and 3 µM HLA, pMHC complex was loaded onto the AKTA Pure system for purification. The fractions containing the correctly refolded pMHC complex were collected and further analyzed.

### Cross-reactivity TCR experiment

Mouse 58-/- T cell hybridoma cells^104^, which expresses mouse CD3 but not TCRab (from David Kranz, University of Illinois) and Chinese hamster ovary (CHO) cells expressing HLA-A2 (generous gift from Dr. James L. Riley, University of Pennsylvania) were cultured in RPMI 1640 medium and DMEM, respectively, supplemented with 10% FBS, sodium pyruvate, non-essential amino acids, glutaMAX-1, penicillin-streptomycin and β-mercaptoethanol. Human variable-mouse constant chimeric gp100(2M)-specific TCR constructs-19LF6 or 16LD6^105^ and human CD8 were retrovirally transduced into the hybridoma cells^106^. Transduced cells were sorted, expanded for 6 days, quantified for TCR and CD3e expression, and prepared for the coculture cytokine assay. 10,000 CHO cells expressing HLA-A2 were loaded with different concentrations of gp100(2M) peptide (IMDQVPFSV), in which anchor position 2 was mutated from original peptide T to M to increase binding affinity and PMEL spliced variant peptide (KTWDQVPFSV) and incubated with 10,000 T cell hybridoma clones expressing human CD8 and 19LF6 or 16LD6 TCR in triplicates for 16 h at 37 °C, 5% CO2. A standard ELISA sandwich was used to quantify cytokine mouse IL-2 production^107^.

### Expansion of spliced PMEL specific TCRs

T cells against the splice peptide were expanded using our previously published methods^108–110^ and previously described protocol for expandingnaïve T cells^111^. Dendritic Cells (DC) from peripheral blood mononuclear cells isolated via Ficoll gradient centrifugation and CD14+ monocytes were isolated using MACS magnetic bead separation, according to the manufacturer’s protocol (Milteniyi, Gaithersburg MD). Briefly, cells were incubated with CD14 beads and MACS buffer for 15 minutes, washed, and applied to pre-wetted LS columns on a magnetic stand, with positive cells eluted and counted. CD14+ monocytes were then cultured in 24 well plates at 1×10⁶ cells/mL in CellGenix (Sartorius CellGenix) media supplemented with glutaMAX (Gibco) in the presence of (GMCSF 800 U/mL) and IL-4 (1200 U/mL). DCs were refreshed with half-media change containing the same cytokines after 2 days, and were pulsed with 1 µg peptide/well for 4 hours after another 2 days. They are then matured with media containing GMCSF (800 U/mL), IL-4 (1200 U/mL), TNFα (10 ng/mL), IL-1β (10 ng/mL), IL-6 (100 ng/mL), PGE2 (1 µg/mL), and IFNγ (100 U/mL). The next day, DCs were pulsed with another 1 µg peptide/well and used to stimulate T cells in the presence of 30 ng/mL IL-21. After 2 days of co-culture, media was refreshed with 10 ng/mL IL-7 and 5 ng/mL IL-15. Cells were grown for 9-12 days with media changes/splits as necessary. After 9-12 days, cells were restimulated with DC (generated as above) in the presence of 5 ng/mL IL-7 and 5 ng/mL IL-15. Media was refreshed after 3 days with 75 U/mL IL-2. After another week, cells were restimulated with DC (generated as above) in the presence of 5 ng/mL IL-15. Media was refreshed after 3 days with 100 U/mL IL-2. Cells were then harvested and used for downstream functional assays.

### Spliced PMEL peptide TCR functional assays

Stably eGFP-transduced target cells were plated in complete medium with or without 50 U/mL IFN-γ for 24 hours. Cells were then rinsed twice with PBS, detached, counted, and replated at 15,000 cells/well (SKMEL-5 and A375) or 30,000 cells/well (SW620) and cultured overnight to allow cells to adhere. Cryopreserved sPMEL-expanded T-cells were thawed and cultured overnight in complete RPMI-1640 supplemented with 5 ng/mL human IL-7 and IL-15 (PeproTech). The following day, T-cells were counted, resuspended in complete medium, and added to target cells at an effector-to-target (E:T) ratio of 2:1 for activation assays. Where indicated, sPMEL peptides were added at a final concentration of 2 μM. Activation Assay After 24 hours, T-cells were harvested and stained for the following markers: CD3 (AF488, clone UCHT1 – BioLegend), CD4 (APC-Fire750, clone RPA-T4 – BioLegend), sPMEL-HLA-A02:01 tetramer (KTWDQVPFSV, APC - produced in-house), canonical KTWGQYWQV-HLA-A02:01 tetramer, PE - produced in-house), CD69 (BV711, clone FN50 – BioLegend), CD137 (PE-Cy7, clone 4B4-1 – BioLegend), and CD25 (BV421, clone BC96 – BioLegend). Samples were acquired on a Symphony A5 flow cytometer (BD Biosciences) and analyzed using FlowJo software (FlowStar).

### Personalized cancer vaccine analysis in melanoma

One melanoma patient with the highest tumor mutational burden (TMB) was selected from the Hartwig Medical Foundation (HMF) cohort. Whole-genome sequencing (WGS) data from the patient’s tumor lesion and matched normal tissue were processed using the HMF Purple pipeline (https://github.com/hartwigmedical/hmftools/blob/master/purple/README.md) to identify somatic mutations, and Variant Effect Predictor (VEP) was applied to predict the protein coding effect. RNA-Seq data were analyzed using the ImmunoVerse pipeline to determine the patient’s HLA alleles and expressed somatic mutations, thereby constructing a comprehensive patient-specific antigen search space encompassing all mutation-derived neoantigens. All predicted neoantigens were queried against 267 publicly available melanoma immunopeptidomes using Tesorai Search, applying a consistent false discovery rate (FDR) threshold of 5%. For each expressed somatic mutation-derived neoantigen, observed MS2 spectra were compared to *in silico* predicted spectra generated by MS2PIP (Immuno-HCD model). Spectral match quality was evaluated using both Pearson correlation and cosine similarity. Binding affinity was predicted using netMHCpan (version 4.1), and immunogenicity was assessed using DeepImmuno web server (https://deepimmuno.research.cchmc.org/)^37^.

---

**Supplementary Figure 1.**
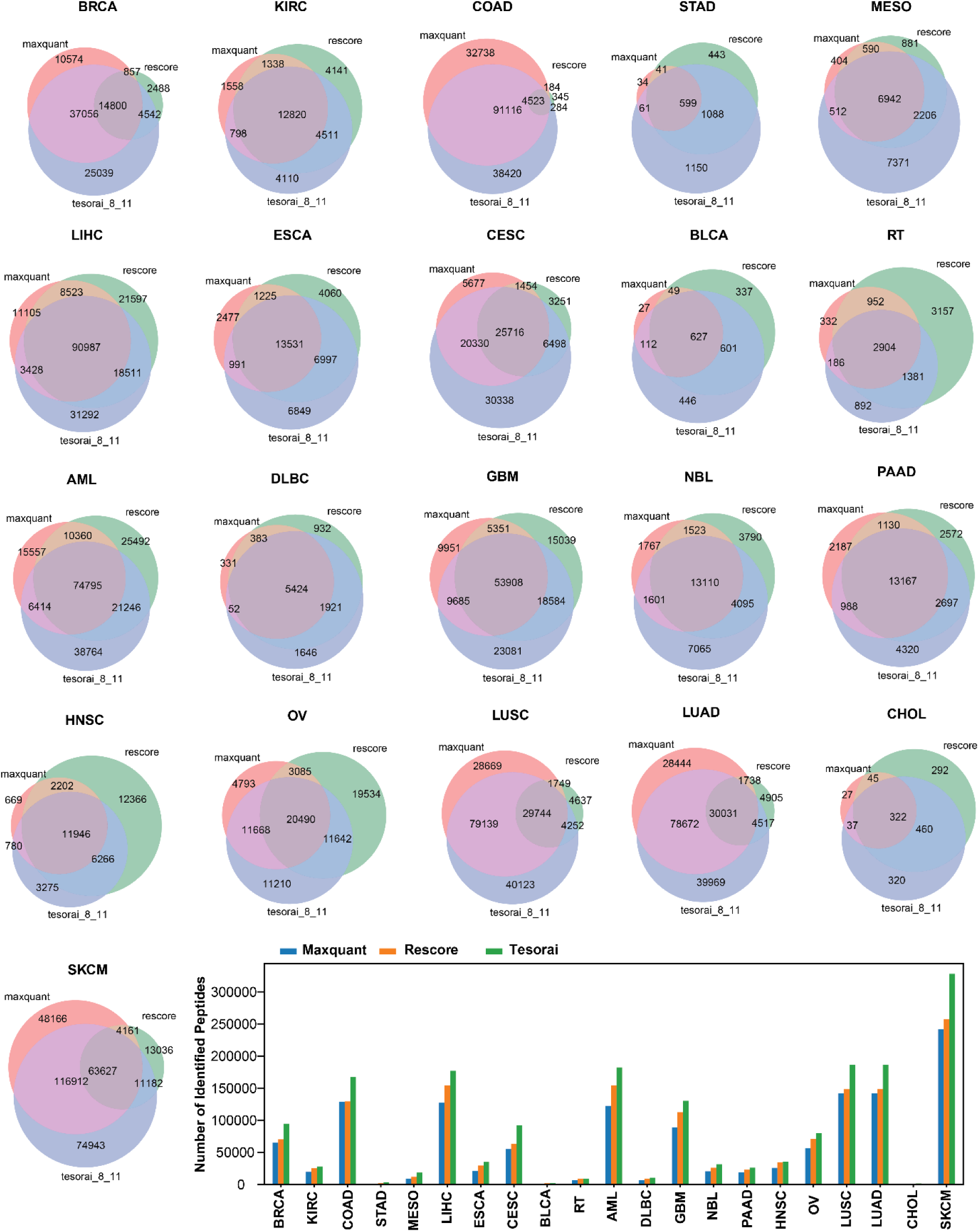
Comparison of identified peptides across MaxQuant, MS2Rescore, and Tesorai Search results across 21 cancers. Venn diagrams illustrate the overlap and unique peptides identified by each algorithm across different cancer types. Barplot summarizes the total number of identified peptides per cancer type, highlighting the relative increase achieved by MS2Rescore and Tesorai Search compared to MaxQuant as baseline.

**Supplementary Figure 2.**
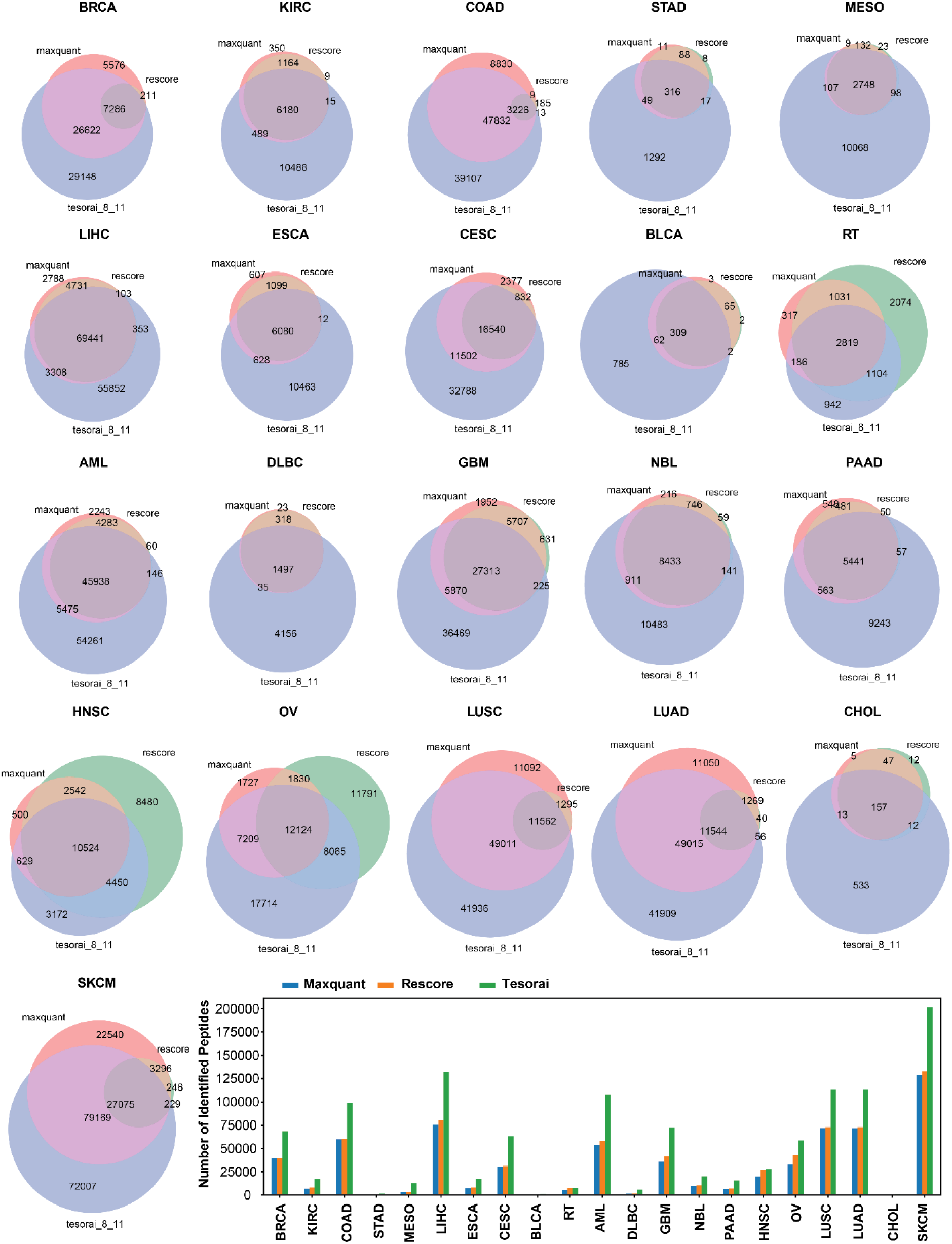
Comparison of identified peptides with high-confidence across MaxQuant, MS2Rescore, and Tesorai search results. Venn diagrams illustrate the overlap and unique peptides identified by each algorithm across different cancer types. Andromeda score > 70 and Tesorai Score > 5 were used to determine high-confidence peptides after consulting with the original developers. Barplot summarizes the total number of identified peptides per cancer type, highlighting the relative increase achieved by MS2Rescore and Tesorai Search compared to MaxQuant as baseline.

**Supplementary Figure 3.**
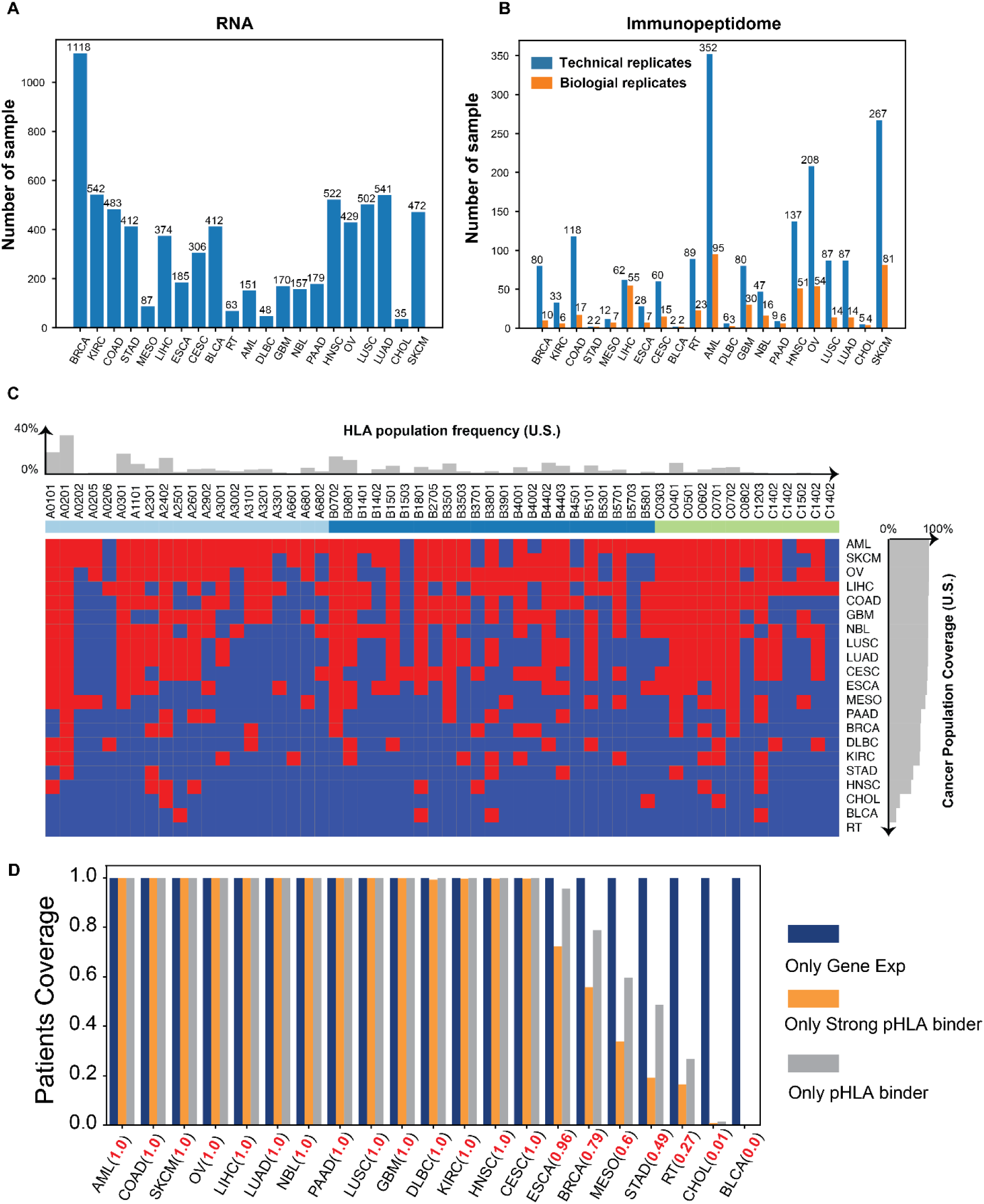
Key statistics associated with public RNA-Seq and immunopeptidome datasets. (A) Number of RNA-Seq datasets for each tumor across TCGA and TARGET. (B) Number of immunopeptidomic datasets for each tumor collected throughout public repositories, number of technical and biological replicates were annotated (C) HLA coverage across cancer types. Bar plots display common HLA alleles (with U.S. population frequency >1%) and their corresponding population coverage, grouped by HLA-A, HLA-B, and HLA-C categories. Red and blue indicate the presence and absence, respectively, of the HLA allele in each cancer cohort. (D) Patient populations potentially eligible for PC-CAR therapy, as evaluated based on gene expression alone and further stratified by tumor-specific HLA restrictions. Both strong and weak peptide-HLA binders were considered in the analysis.

**Supplementary Figure 4.**
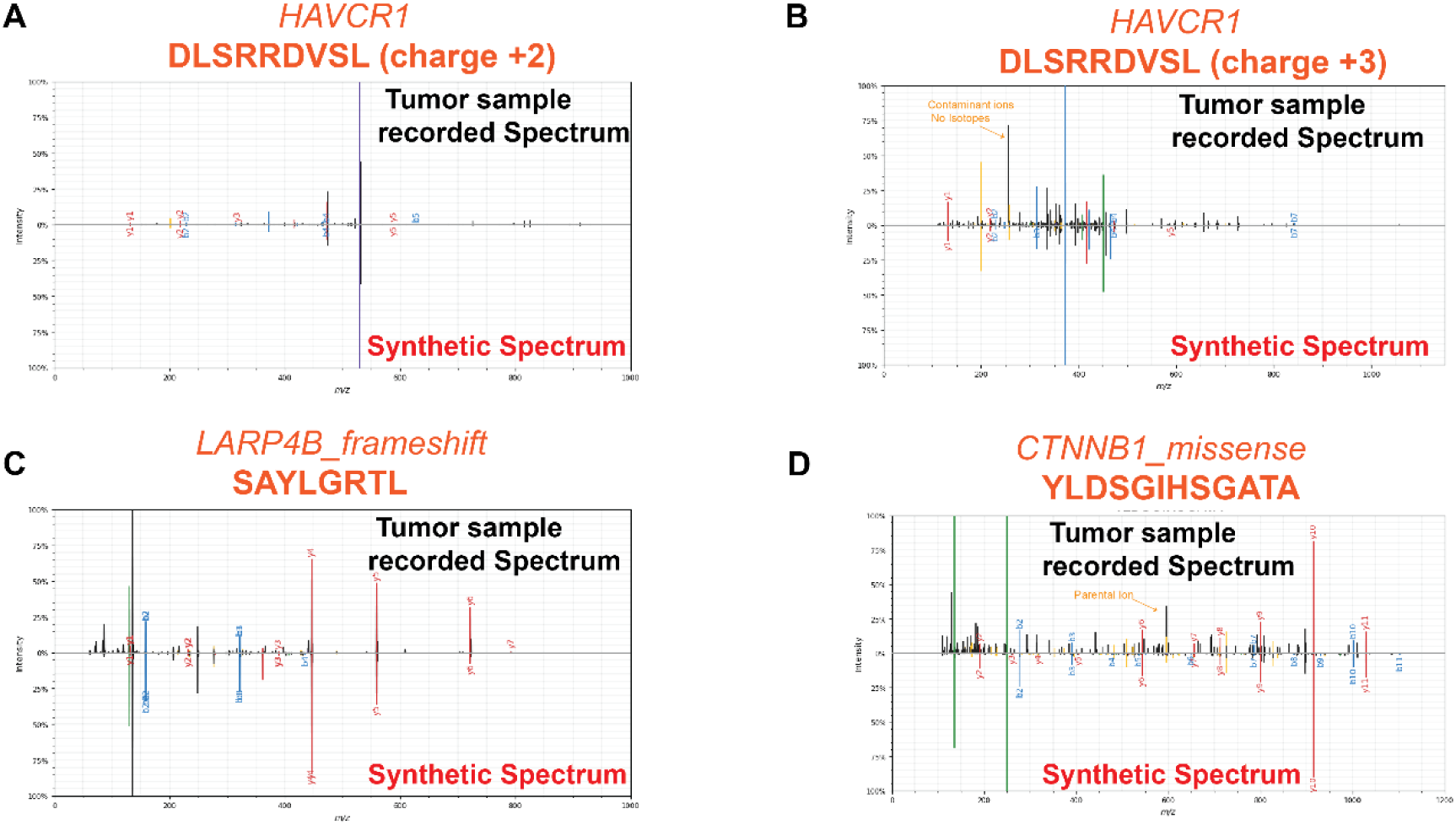
Spike-in mass spectrometry validation of HAVCR1 peptides and mutation-derived neoantigens. (A–B) Mirror plots for HAVCR1 peptides at charge states +2 and +3, respectively, representing the highest-scoring spectra identified by MaxQuant and Tesorai Search, compared to corresponding synthetic spectrum. (C) Mirror plot for the frameshift mutation–derived neoantigen SAYLGRTL from LARP4B. (D) Mirror plot for the missense mutation–derived neoantigen YLDSGIHSGATA from CTNNB1.

**Supplementary Figure 5.**
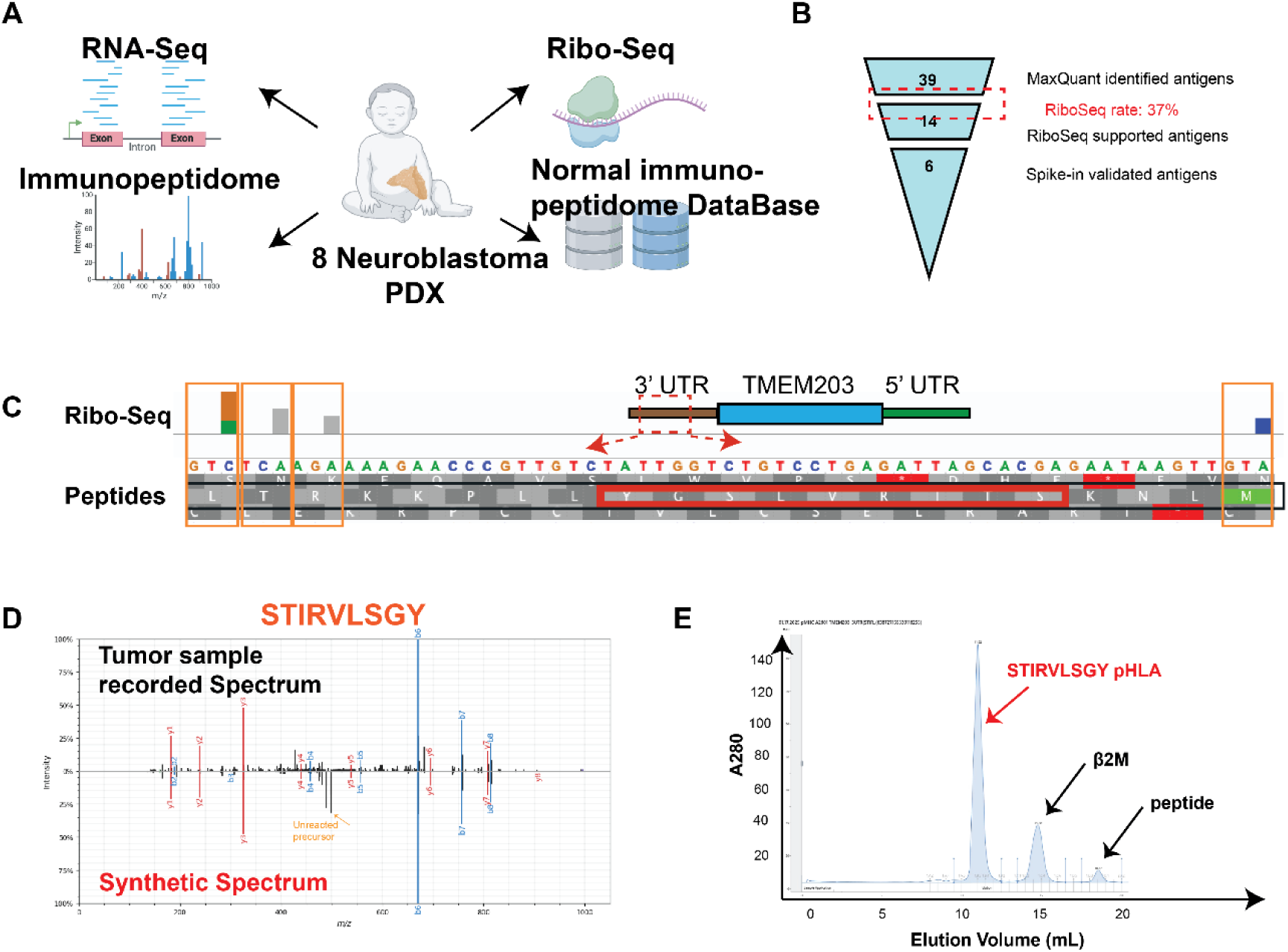
Cryptic ORF antigen validation workflow and TMEM203 3’ UTR derived STIRVLSGY antigen. (A) workflow of assessing translational activity of cryptic ORFs using Ribo-Seq in matched PDX neuroblastoma models (B) funnel plot of validated cryptic ORF antigen in matched Neuroblastoma PDX model based on Ribo-Seq evidence (C) Ribosome occupancies on 3’ UTR of TMEM203 from Ribo-Seq, the STIRVLSGY peptide (residing on negative strand) was highlighted in red box and the Ribo-Seq confirmed codons were highlighted in orange box (D) Mirror plot of synthetic peptide and original spectra to confirm the Mass Spectrometry (MS) identification of TMEM203 cryptic ORF antigen (E) Size exclusion chromatography after refolding the STIRVLSGY-HLA-A*26:01 complex.

**Supplementary Figure 6.**
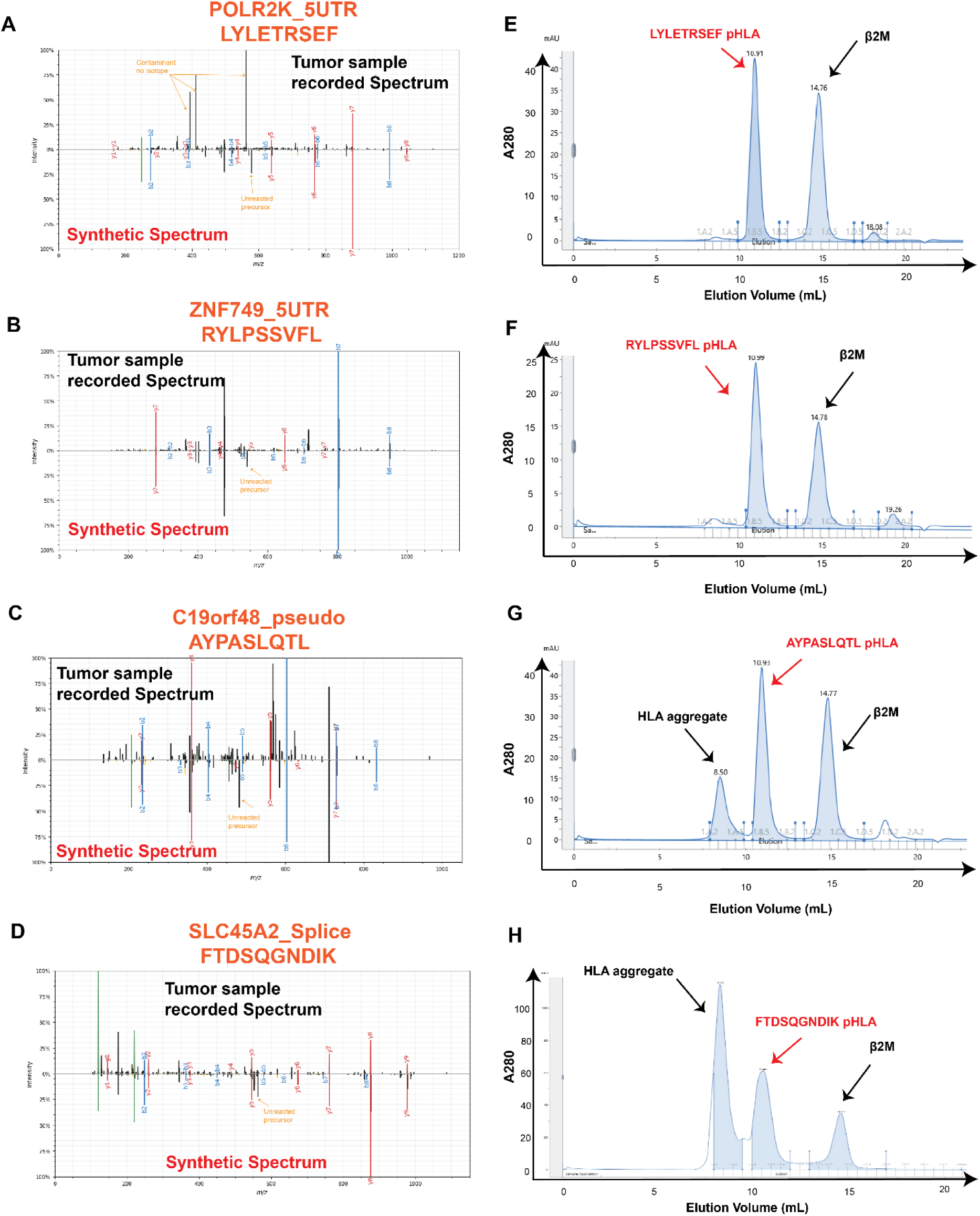
Validation of additional Cryptic ORF and splicing antigens. (A-D) Mirror plots of original spectrum from the tumor samples and the synthetic peptide spectra to validate the MS identification for POLR2K_5UTR, ZNF749_5UTR, C19orf48_pseudogene, SLC45A2_spliced (top to bottom). (E-H) Size Exclusion Chromatography after refolding pHLA for LYLETRSEF-A2402 complex, RYLPSSVFL-A2402 complex, AYPASLQTL-A2402, FTDSQGNDIK-A1101 complex (top to bottom).

**Supplementary Figure 7.**
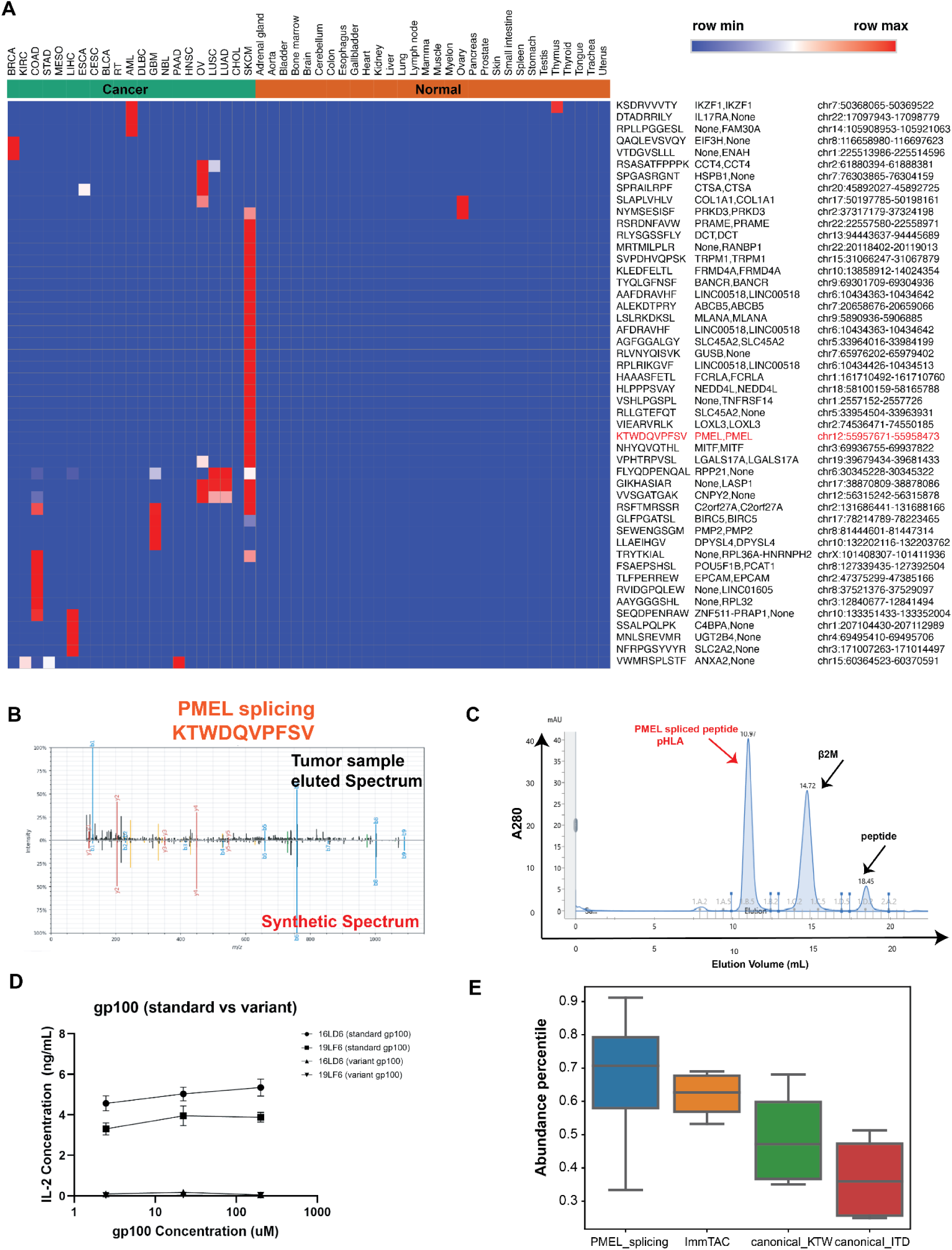
Splicing antigen landscapes across tumor types. (A) High-confidence and frequently detected splicing antigen across 21 cancers, annotated by the parental genes of the two splicing sites (splicing acceptor side and splicing donor site, None denotes the splicing sites are unannotated novel splicing sites) and chromosomal coordinates (B) Validation of PMEL spliced peptide using synthetic peptide MS/MS (C) Size exclusion chromatography after refolding PMEL spliced peptide and HLA-A*02:01 allele (D) TCR-T co-culture ELISA assay using specific TCR for canonical ITD antigens toward both canonical ITD and spliced variant (E) Comparison of peptide abundance (expressed as percentile of abundance) amongst four PMEL antigens.

**Supplementary Figure 8.**
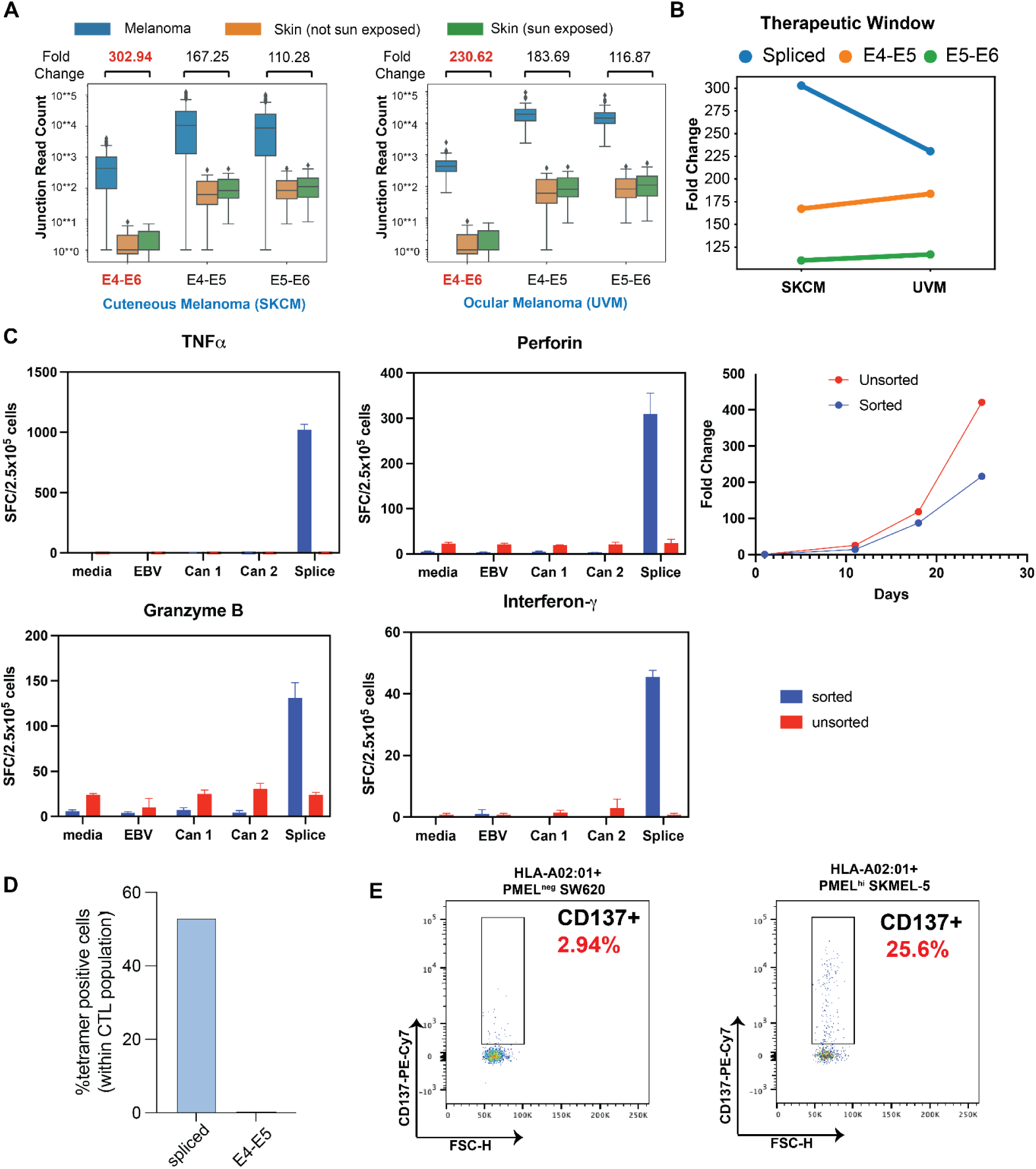
Therapeutic windows of spliced PMEL (sPMEL) and enrichment of sPMEL-specific TCRs. (A) Therapeutic windows on RNA level between spliced and canonical junctions in both cutaneous melanoma and ocular melanoma. The fold change is derived between melanoma and sun exposed skin (B) The changes of therapeutic windows between cutaneous melanoma and ocular melanoma for PMEL spliced antigens and two canonical counterparts. (C) sPMEL-tetramer presorted T cells from a healthy donor exhibit specific expansion in response to sPMEL antigen, as compared to control tetramers loaded with the Can1 peptide (E4–E5 junction), Can2 peptide (E5–E6 junction), and a positive control peptide from EBV (CLGGLLTMV). Antigen-specific responses were assessed by measuring secretion levels of TNFα, perforin, granzyme B, and interferon-γ, as well as fold changes in cell numbers over the course of expansion. (D) Binding to spliced-derived peptide as well as canonical (KTWGQYWQV) peptide is quantified within the CTL population. (E) Representative flow plots showing CD137 expression within CTL population without stimulation or 20h after co-culture with HLA-A02:01+Pmel^high^ SKMEL-5 melanoma cell line.

**Supplementary Figure 9.**
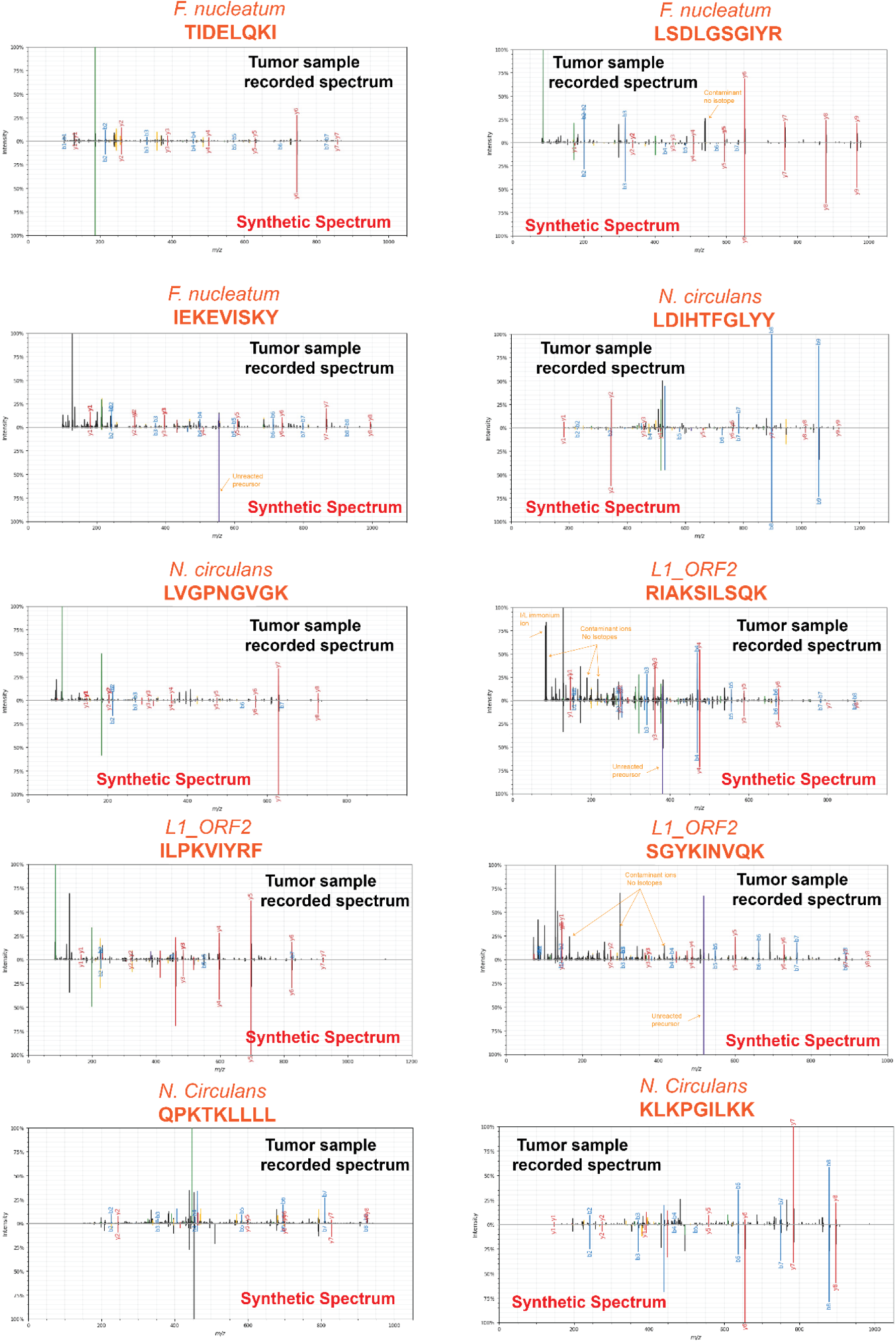
Validation of pathogen and TE derived antigens. Mirror plots comparing the original MS/MS spectra from tumor samples with the corresponding synthetic peptide spectra to validate the identification of nine pathogen-derived peptides. These include peptides from *Fusobacterium nucleatum* (TIDELQKI, LSDLGSGIYR, IEKEVISKY), *Niallia circulans* (LDIHTFGLYY, LVGPNGVGK), and L1_ORF2 (RIAKSILSQK, ILPKVIYRF, SGYKINVQK). Two additional peptides derived from *Niallia circulans* (QPKTKLLLL and KLKPGILKK), which exhibit high spectral similarity, were acquired in the tumor sample using Wideband Activation mode that is only available for the linear trap quadrupole (LTQ) Orbitrap XL mass spectrometer. This instrument is not available for the analysis of the synthetic peptide, instead, Multistage Activation was used for these two peptides, targeting also the M-17 ion for fragmentation.

**Supplemental Figure 10.**
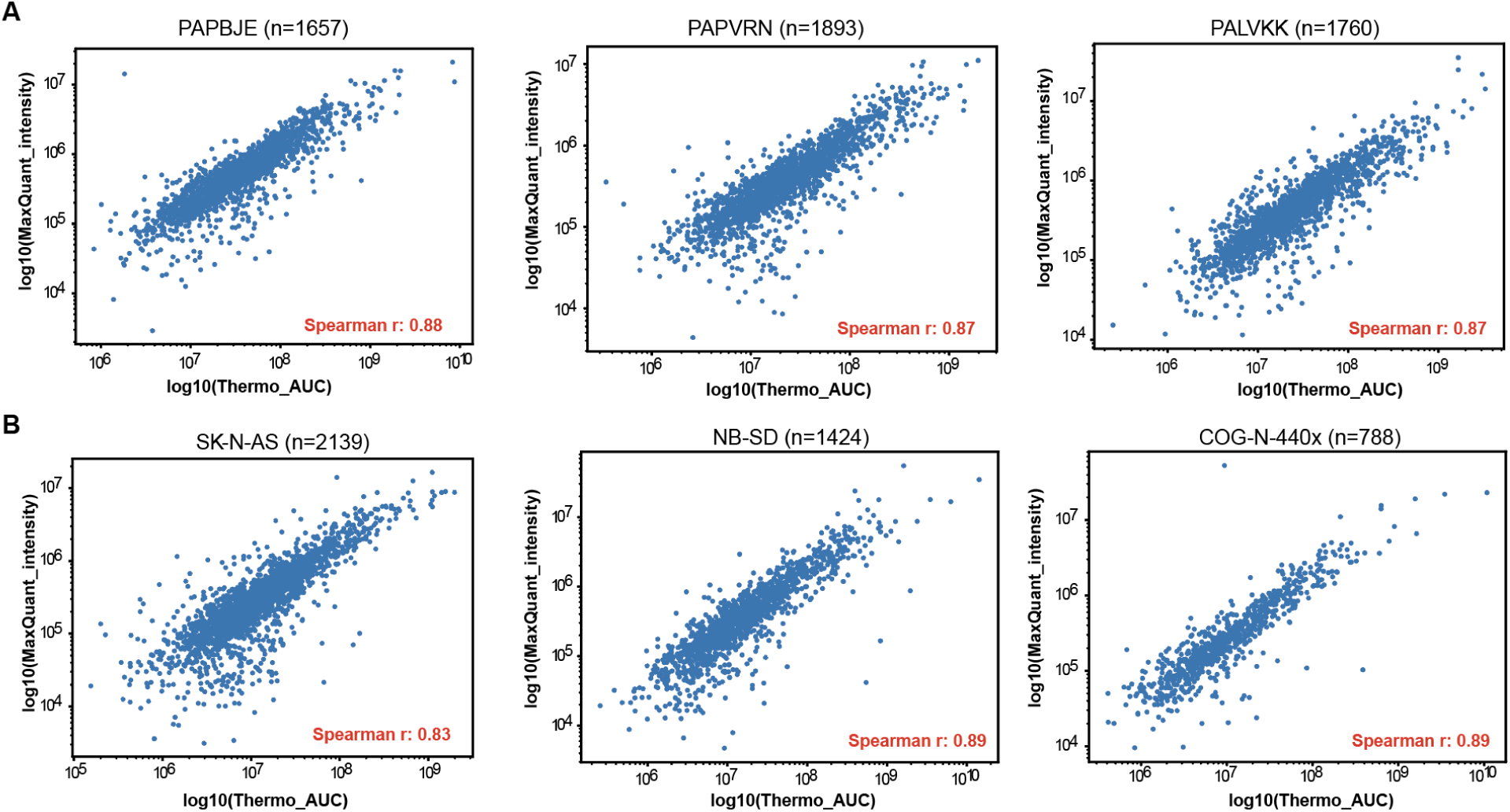
Correlation between maximum precursor intensity reported by MaxQuant and Area Under the Curve (AUC) reported by Thermo proprietary software (ProteomeDiscoverer). (A) Spearman correlation of intensity values for commonly detected peptides in three neuroblastoma primary tumors (PAPBJE, PAPVRN, PALVKK) with the highest number of overlapping identifications. (B) Spearman correlation of intensity values for commonly detected peptides in three neuroblastoma cell lines (SK-N-AS, NB-SB, COG-N-440x) with the highest overlap.

## Notes

https://github.com/frankligy/pan_cancer_intracellular_antigen_atlas

https://www.immuno-verse.com/

https://frankligy.github.io/pan_cancer_intracellular_antigen_atlas/notes/deployed_pipeline_latest.pdf)

### Summary of Updates

1. Enhanced sensitivity for antigen discovery through a novel AI-powered model (Tesorai), enabling broader identification of tumor-specific antigens. 2. Experimental validation of ex vivo expansion of T cells specific to the spliced PMEL antigen, demonstrating the functional relevance of reported tumor-specific antigens. 3. Comprehensive web portal and analysis pipeline developed for user-friendly access and reproducible workflows. 4. Expanded discovery and validation of mutation-derived neoantigens across diverse datasets. 5. New application in personalized medicine by leveraging public immunopeptidome datasets for individualized antigen design

